# Loss of prohibitin 2 in Schwann cells dysregulates key transcription factors controlling developmental myelination

**DOI:** 10.1101/2024.03.20.585915

**Authors:** Emma R. Wilson, Gustavo Della-Flora Nunes, Shichen Shen, Seth Moore, Joseph Gawron, Jessica Maxwell, Umair Syed, Edward Hurley, Meghana Lanka, Jun Qu, Laurent Desaubry, Lawrence Wrabetz, Yannick Poitelon, M. Laura Feltri

## Abstract

Schwann cells are critical for the proper development and function of the peripheral nervous system, where they form a mutually beneficial relationship with axons. Past studies have highlighted that a pair of proteins called the prohibitins play major roles in Schwann cell biology. Prohibitins are ubiquitously expressed and versatile proteins. We have previously shown that while prohibitins play a crucial role in Schwann cell mitochondria for long-term myelin maintenance and axon health, they may also be present at the Schwann cell-axon interface during development. Here, we expand on this work, showing that drug-mediated modulation of prohibitins *in vitro* disrupts myelination and confirming that Schwann cell-specific ablation of prohibitin 2 (*Phb2*) *in vivo* results in early and severe defects in peripheral nerve development. Using a proteomic approach *in vitro*, we identify a pool of candidate PHB2 interactors that change their interaction with PHB2 depending on the presence of axonal signals. Furthermore, we show *in vivo* that loss of *Phb2* in mouse Schwann cells causes ineffective proliferation and dysregulation of transcription factors EGR2 (KROX20), POU3F1 (OCT6) and POU3F2 (BRN2) that are necessary for proper Schwann cell maturation. Schwann cell-specific deletion of *Jun*, a transcription factor associated with negative regulation of myelination, confers partial rescue of the development defect seen in mice lacking Schwann cell *Phb2*. This work develops our understanding of Schwann cell biology, revealing that *Phb2* may directly or indirectly modulate the timely expression of transcription factors necessary for proper peripheral nervous system development, and proposing candidates that may play a role in PHB2-mediated integration of axon signals in the Schwann cell.

## Introduction

Within both the central and peripheral nervous system (CNS and PNS) axons are enwrapped and supported by glial cells. In addition, the axons of some CNS and PNS neurons are myelinated by concentric layers of lipids and proteins. Schwann cells, the myelinating and main glial cells of the PNS, form an important and symbiotic relationship with neurons, providing trophic and metabolic support (Beirowski, 2019; Boucanova & Chrast, 2020). Thus, Schwann cells are critical for the proper development and maintenance of the peripheral nerve (Wilson et al., 2020) and to promote peripheral nerve regeneration following injury (Jessen & Mirsky, 2016).

From embryonic age 17.5 (E17.5) in mice, Schwann cells select out large diameter axons with which they form a one-to-one relationship and myelinate, in a process known as radial sorting (Feltri et al., 2016). After birth, promyelinating Schwann cells begin to wrap axons in layers of myelin sheath (Feltri et al., 2016). The remaining and typically smaller axons are engulfed by Remak Schwann cells, which do not form myelin. The process of radial sorting is difficult to study *in vivo*. Our lab previously developed an *in vitro* pseudopod system to study the Schwann cell-axon interface (Poitelon & Feltri, 2018), and identified prohibitins (PHBs) as proteins that localize to the Schwann cell membrane in response to neuronal signals (Poitelon et al., 2015).

PHBs (prohibitin 1, PHB1, and prohibitin 2, PHB2) are ubiquitously expressed and evolutionarily conserved proteins. Roles for PHB1 (also known as BAP32) and PHB2 (also known as BAP37 and REA) have been reported in diverse subcellular compartments, including most dominantly in the mitochondria, but also in the nucleus, and at the cell membrane (Osman et al., 2009). We previously showed that PHB1 in Schwann cells contributes to mitochondrial health. By post-natal day 40 (P40), mice lacking PHB1 in Schwann cells (PHB1 Schwann cell knockout, PHB1-SCKO) present with a severe peripheral neuropathy caused by mitochondrial dysfunction, in turn causing activation of the mTORC1 pathway and JUN (also known as c-JUN), which leads to demyelination (Della-Flora Nunes, Wilson, Hurley, et al., 2021; Della-Flora Nunes, Wilson, Marziali, et al., 2021). Conditional ablation of PHB2 from Schwann cells (PHB2 Schwann cell knockout, PHB2-SCKO) results in a severe radial sorting defect in mice that is evident at P20 (Poitelon et al., 2015).

In this study, we investigate the mechanisms by which PHB2 impacts Schwann cells during mammalian peripheral nerve development using *in vitro* models and the previously established PHB2-SCKO mouse model (Poitelon et al., 2015). We confirm genetically and pharmacologically that Schwann cell PHBs play a crucial role in myelination. We also identify Schwann cell proteins that change their interaction with PHB2 depending on the presence or absence of neuronal signals. Our studies reveal that PHB2 knockout from Schwann cells causes dysregulation of transcription factors, namely EGR2 (KROX20), POU3F1 (OCT6) and POU3F2 (BRN2), and that PHB2-lacking Schwann cells in the developing peripheral nerve are not able to readily proliferate. We find that the transcription factor JUN, commonly associated with Schwann cell transdifferentiation to repair Schwann cells and considered a negative regulator of myelination, is elevated in PHB2-SCKO mice and that genetic ablation of *Jun* in Schwann cells confers a partial but significant rescue of the radial sorting defect. Overall, our results contribute to a growing understanding of the holistic formation of the peripheral nerve during development and suggest that while PHBs are found at the Schwann cell-axon interface and play a critical role in maintaining Schwann cell mitochondrial health, they may also contribute, directly or indirectly, to the regulation of Schwann cell transcription factors.

## Materials and Methods

### Animals

All animal experiments in this study followed ethical regulations for animal testing and research and were approved by the Institutional Animal Care and Use Committee (IACUC) of the Roswell Park Cancer Institute and the regulatory authorities at the University at Buffalo. Animals were housed in individually ventilated cages, separated by gender in groups of at most five per cage, kept in a 12 h light/dark cycle, with food and water ad libitum. *Mpz*-Cre (Feltri et al., 1999), *Phb1*-floxed (He et al., 2011), *Phb2*-floxed (Park et al., 2011), and *Jun*-floxed (Behrens et al., 2002) mice were used in this study. Controls contained two, one, or no floxed alleles and were negative for the *Mpz*-Cre gene. Mice were kept in a C57BL/6 background. Littermate controls were used throughout, except for triple transgenic animals, where littermate controls were used whenever possible. No animals were excluded from this study. Genotyping was performed from genomic DNA. For *Mpz*-Cre (450 bp product) the primers used for genotyping were F: 5’ CCACCACCTCTCCATTGCAC 3’, R: 5’ GCTGGCCCAAATGTTGCTGG 3’ and the PCR conditions were 94°C for 5 minutes, 30 x (94°C for 30 seconds, 56°C for 30 seconds, 72°C for 1 minute), 72°C for 10 minutes. For *Phb1* (195 bp for the floxed product, 118 bp for the wildtype product) the primers used for genotyping were F: 5’ TAAGACTGGGTCCTGCCATT 3’, R: 5’ GTGCTTGCATCAGAGTCAGG 3’ and the PCR conditions were 94°C for 10 minutes, 30 x (94°C for 40 seconds, 57°C for 40 seconds, 72°C for 40 seconds), 72°C for 10 minutes. For *Phb2* (647 bp for the floxed product, 277 bp for the wildtype product) the primers used for genotyping were F: 5’ ATGAGACCAAGGCGAATG 3’, R: 5’ GCTGTCAGATTCAAGGTTGC 3’ and the PCR conditions were 95°C for 2 minutes, 30 x (95°C for 30 seconds, 58°C for 30 seconds, 72°C for 2 minutes), 72°C for 7 minutes. For *Jun* (350 bp for the floxed product, 300 bp for the wildtype product) the primers used for genotyping were F: 5’ CCGCTAGCACTCACGTTGGTAGGC 3’, R: 5’ CTCATACCAGTTCGCACAGGCGGC 3’ and the PCR conditions were 94°C for 3 minutes, 5 x (94°C for 30 seconds, 67°C (-1°C per cycle) for 30 seconds, 72°C for 1 minute), 25 x (94°C for 30 seconds, 62°C for 30 seconds, 72°C for 1 minute), 72°C for 5 minutes.

### Morphological assessments

Sciatic nerves were fixed in 2% glutaraldehyde and stored at 4°C. Tissue was post-fixed in 1% osmium tetroxide, dehydrated by stepwise sequential incubation in ethanol of increasing concentration, and embedded in Epon resin using propylene oxide as a transition solvent. 1 μm thick sections were stained with 2% toluidine blue and images were acquired using the ×100 objective of a Leica DM6000 microscope and the Q-Capture Pro V7.0.4324.5 software (QImaging, Inc.). PTGui software v.10 (New House Internet Services BV) was used to stitch together the images to reconstruct the complete cross-section of the nerve and the number of myelinated fibers, amyelinated fibers, and improperly sorted bundles was quantified from the whole nerve using the cell counter plugin from ImageJ2 Fiji v2.14.0/1.54f (Rueden et al., 2017; Schindelin et al., 2012). Ultrathin sections (80-85 nm) were stained with uranyl acetate and lead citrate to be examined using electron microscopy at ×2900 magnification. The number of axons in unsorted bundles were quantified from 23-35 random images, using the cell counter plugin from ImageJ2 Fiji software.

### Myelinating co-cultures

Purified DRG neuron cultures were prepared from E15.5 rat embryos, according to the protocol outlined in (Taveggia & Bolino, 2018). 12 mm acid-washed glass coverslips were coated with collagen (50 µg/mL in a 0.1% glacial acetic acid solution in water) and matrigel (1:10 in ice cold DMEM, brief coating). On day 9 400,000 primary rat Schwann cells isolated from postnatal day 3-7 rat sciatic nerves (using the Brockes’ method and described by our group previously (Brockes et al., 1979; Della-Flora Nunes, Wilson, Marziali, et al., 2021)) were plated onto purified DRG neuron cultures. From day 10, the co-cultures were grown in myelinating cell culture medium (C medium) for 11 further days. The PHB modulating compounds were solubilized in DMSO to make 10 mM stock solutions and then diluted in the myelinating cell culture medium (C medium) to the concentrations stated in the figures and included in all subsequent media changes. DMSO was used as a vehicle control. On day 20, cultures were fixed in 4% paraformaldehyde for 20 minutes, permeabilized for 10 minutes in methanol and blocked for 1 hour at room temperature in 1x PBS, 20% fetal bovine serum, 2% bovine serum albumin, 0.1% Triton X-100. Myelin internodes were detected by immunostaining for β-tubulin III (TUBB3) and myelin basic protein (MBP) (Table 1) with corresponding fluorophore-conjugated secondary antibodies. Nuclei were identified using DAPI. Coverslips were mounted in Vectashield (Vector Laboratories). 3-6 fields of view were taken using the ×20 objective of a Zeiss apotome microscope (in standard fluorescence mode) and AxioVision 4.8 software. Internode numbers were quantified using an automated routine for ImageJ2 Fiji. Briefly, images were loaded in the software and the scale set. Images were then filtered using a Gaussian blur filter and image subtraction was applied to reduce background noise. Myelin internodes were selected using thresholding and their length (Feret’s diameter) was measured automatically. Objects smaller than 40 µm^2^ or with circulatory values higher than 0.3 were excluded from the analysis since they often represented non-specific staining. Overlapping internodes, which could not be correctly individualized by our protocol, were also excluded. For each condition, the number of internodes is expressed as a percentage of the number of internodes in untreated controls for that experimental replicate. Our semi-automated analysis was able to produce results in line with a manual analysis approach (Supp. Fig. 1A-C).

**Table 1:** Primary antibodies used. WB = western blot. IHC = immunohistochemistry.

| **Antibody** | **Source species** | **Dilution**<br>**(experiment)** | **Company (catalog**<br>**#)** |
| --- | --- | --- | --- |
| MBP | Rat | 1:5 (IHC) | Hybridoma |
| TUBB3 | Rabbit | 1:1000 (IHC) | Biolegend (802001) |
| PHB2 | Rabbit | 1:400 (WB) | Atlas (HPA039874) |
| ERK1/2 | Rabbit | 1:750 (WB) | Cell Signaling (9102) |
| phospho-ERK1/2<br>(Thr202/Tyr204) | Rabbit | 1:750 (WB) | Cell Signaling (9101) |
| β-tubulin | Rabbit | 1:1000 (WB) | Novus Biologicals<br>(NB600-936) |
| AKT | Rabbit | 1:750 (WB) | Cell Signaling (9272) |
| phospho-AKT<br>(Ser473) | Rabbit | 1:750 (WB) | Cell Signaling (9271) |
| S6 ribosomal protein | Rabbit | 1:750 (WB) | Cell Signaling (2217) |
| phospho-S6<br>ribosomal protein<br>(Ser235/236) | Rabbit | 1:750 (WB) | Cell Signaling (4858) |
| 4E-BP1 | Rabbit | 1:750 (WB) | Cell Signaling (9644) |
| phospho-4E-BP1<br>(Thr37/46) | Rabbit | 1:750 (WB) | Cell Signaling (2855) |
| EGR2/KROX20 | Rabbit | 1:2000 (WB)<br>1:8000 (IHC) | gift from Dies Meijer |
| POU3F1/OCT6 | Rabbit | 1:2000 (WB)<br>1:500 (IHC) | gift from Dies Meijer |
| POU3F2/BRN2 | Rabbit | 1:1000 (WB) | Cell Signaling<br>(12137) |
| JUN | Rabbit | 1:750 (WB)<br>1:200 (IHC) | Cell Signaling (9165) |
| SOX10 | Rabbit | 1:1000 (WB)<br>1:500 (IHC) | Cell Signaling<br>(89356) |
| MKI67 | Rat | 1:300 (IHC) | Invitrogen<br>(14-5698-80) |
| CANX | Rabbit | 1:3000 (WB) | Sigma-Aldrich<br>(C4731) |

### TurboID

For turboID experiments, purified DRG neuron cultures and primary rat Schwann cells were prepared as per the myelinating co-cultures, in 6 well plates. Primary rat Schwann cells were transfected with either con-turboID or PHB2-turboID plasmids using lipofectamine 3000 according to the manufacturer’s instructions. After 8 days of culturing DRG neurons, transfected rat Schwann cells were seeded onto the neurons in medium containing 50 µM biotin. After 2 hours, cultures were washed in PBS and protein harvested in lysis buffer (150 mM sodium chloride, 10 mM sodium fluoride, 1% IGEPAL CA-630, 1% protease inhibitor cocktail (Sigma-Aldrich P8340), 1% phosphatase inhibitor cocktail 2 (Sigma-Aldrich P5726), 1% phosphatase inhibitor cocktail 3 (Sigma-Aldrich P0044), 1X PBS, pH 7.6). Protein was quantified using the BCA protein assay (ThermoFisher Scientific) according to the manufacturer’s instructions. Biotinylated proteins were purified using Dynabeads MyOne Streptavidin T1 (Invitrogen). Briefly, Dynabeads were washed three times in PBS then incubated with 100 µg (Schwann cell alone condition) or 300 µg (Schwann cells with DRG neurons condition) for 1 hour at room temperature while rotating. Dynabeads were then washed four times in PBS and half of the Dynabeads used for liquid chromatography-mass spectrometry (LC-MS) analysis and half used for assessment via western blot. Western blot of turboID samples was carried out as described below and blots were probed with an antibody recognizing PHB2 (Table 1) or HRP-conjugated streptavidin to visualize biotinylated proteins.

### Liquid chromatography-mass spectrometry (LC-MS) and data analysis

Following on-bead digestion, LC-MS was performed as described previously (Kalem et al., 2022). Within each experimental condition and for each independent experiment, the total ion chromatograph (TIC) intensity for each protein was calculated relative to the total TIC intensity for all the biotinylated proteins pulled down by streptavidin-affinity purification (AP), so that we compare the percentage contribution of each protein to the total biotinylated pool. Next, we calculated the ratio of the relative TIC intensity for PHB2-TurboID interactors when in the presence of biotin to in the absence and eliminated proteins with a ratio of lower than 1 as non-specific. When examining the increased interactors, we removed non-specifics by comparing group 4 (Schwann cells with neurons plated with biotin) to group 6 (Schwann cells with neurons plated without biotin). When examining decreased interactors, we removed non-specifics by comparing group 3 (Schwann cells alone with biotin) to group 5 (Schwann cells alone without biotin). This was to retain candidates in our analysis workflow which were present only when Schwann cells were plated alone. Finally, we calculated the ratio of TIC intensities of PHB2-turboID interactors when Schwann cells were plated with DRG neurons to when they were plated alone. Proteins with a ratio of 1 or more on three independent experimental repeats were categorized as increasing PHB2 interactors in the presence of neuronal signals. In contrast, proteins with a ratio of less than 1 on all three independent experimental repeats were categorized as decreasing their interaction with PHB2 in the presence of neuronal signals. Uncharacterized proteins were removed.

### Western blot

Sciatic nerves were dissected and from animals aged post-natal day 20 (P20) or more, the epineurium was removed (removal of the epineurium was not possible from the nerves of younger animals). For triple transgenic animals at P5, sciatic nerves were combined with the brachial plexuses. Tissues were immediately snap-frozen in liquid nitrogen and stored at -80°C until processing. Frozen nerves were pulverized using a pestle and mortar and resuspended in lysis buffer (50 mM Tris pH 7.4, 150 mM sodium chloride, 1% IGEPAL CA-630, 1 mM EDTA, 1 mM EGTA, 1 mM sodium orthovanadate, 0.1% SDS, 1 mM sodium fluoride, protease inhibitor cocktail (Sigma-Aldrich P8340), phosphatase inhibitor cocktail 2 (Sigma-Aldrich P5726) and phosphatase inhibitor cocktail 3 (Sigma-Aldrich P0044)). The samples were placed in a cell agitator for 10 min at 4°C and subsequently subjected to three 20-second cycles of sonication (70% power) in a water sonicator. Following centrifugation at 16,000 x *g* for 15 min at 4°C, the supernatant was collected and the protein concentration was determined by BCA protein assay (Thermo Fisher Scientific) according to the manufacturer’s instructions. Equal amounts of protein per sample were diluted 3:1 in 4X reducing Laemmli SDS sample buffer and denatured for 5 min at 100°C. Proteins were resolved by SDS-PAGE and transferred onto a PVDF membrane. Membranes were blocked with 5% BSA in TBS-T (1× TBS + 0.1% Tween-20) and incubated overnight at 4°C with primary antibody, in 1-5% BSA in TBS-T. The primary antibodies used for western blots are listed in Table 1. Membranes were washed three times in 1x TBS-T and incubated with HRP-conjugated secondary antibody (Novus Biologicals NB7185) for 1 h at room temperature. Following three further washes in TBS-T, membranes were developed using Pierce ECL (Thermo Fisher Scientific) or Amersham ECL Select (Cytiva) western blotting detection reagent and imaged using a ChemiDoc XRS+ system. Quantifications were carried out in the Image lab 6.0 software (BioRad). Band intensity was normalized to the loading controls β-tubulin or calnexin and subsequently expressed relative to the average of control animals or to a common control band. Uncropped blots are presented in Supp. Fig. 7 and 8.

### Immunohistochemistry

Dissected sciatic nerves were snap-frozen in OCT. A cryostat was used to cut 10 µm thick longitudinal sections which were mounted on slides and post-fixed for 10 minutes in ice cold 4% PFA. Sections were blocked for 1 hour at room temperature in 1x PBS, 20% fetal bovine serum, 2% bovine serum albumin, 1% Triton X-100. Slides were incubated overnight in a humidity chamber at 4°C with primary antibodies (Table 1) and following three PBS washes incubated for 1 hour at room temperature with appropriate fluorophore-conjugated secondary antibodies. Nuclei were identified using DAPI. After PBS washes, slides were mounted in Vectashield (Vector Laboratories). Images were acquired using the ×20 objective on a confocal microscope Leica SP5II running the LAS AF 2.7.9723.3 software (Leica) and analyzed blind to genotypes using the cell counter plugin of the ImageJ2 Fiji software.

### TUNEL

For identification of apoptotic nuclei by terminal deoxynucleotidyl transferase dUTP nick end labeling (TUNEL) dissected sciatic nerves were frozen, sectioned, and post-fixed as for immunohistochemistry. Sections were permeabilized for 1 minute in ice cold methanol. Sections were incubated for 15 minutes at room temperature in TdT buffer (30 mM Tris, 2 mM cobalt chloride, 140 mM sodium cacodylate). Sections were then incubated with terminal transferase (TdT, 0.8 units/ µL) with biotin-16-dUTP in TdT buffer for 1 hour at 37°C. Following a PBS wash, the TdT reaction was blocked with buffer containing 300 mM sodium chloride and 40 mM sodium citrate monobasic. Sections were then blocked for 1 hour at room temperature in 2% bovine serum albumin and then incubated with streptavidin-rhodamine (1:400 in 2% BSA) for 1 hour at room temperature. Nuclei were identified using DAPI. After PBS washes, slides were mounted in Vectashield (Vector Laboratories). Images were acquired using the ×20 objective on a confocal microscope Leica SP5II running the LAS AF 2.7.9723.3 software (Leica) and analyzed blind to genotypes using the cell counter plugin from ImageJ2 Fiji software.

### Quantitative reverse transcriptase PCR (qRT-PCR)

Dissected sciatic nerves (epineurium removed) were snap-frozen in liquid nitrogen. Frozen nerves were pulverized with a pestle and mortar and TRIzol (Invitrogen) used to extract RNA as per the manufacturer’s instructions. RNA was reverse transcribed into cDNA using Superscript III Reverse Transcriptase (Invitrogen) according to the manufacturer’s instructions, using both oligo(dT)_20_ and random primers. For quantitative PCR the FastStart Universal Probe system (Roche) was used. Data were analyzed using the 2^−ΔΔCt^ method relative to the housekeeping *Actb* gene and the average ΔCt of control animals. Fold change was then normalized to the average fold change in control animals. The primers and probes used were as follows: *Phb2*: forward 5’ CCATTGTTAATGAGGTGCTCAA 3’, reverse 5’ CTTCGGATCAACAGGGACA 3’, probe 47, *Phb1*: forward 5’ GGGTCCTGCCTTCTATCACC 3’, reverse 5’ TCAATTCTCCAGCATCGAATC 3’, probe 77, *Actb*: forward 5’ AAGGCCAACCGTGAAAAGAT 3’, reverse GTGGTACGACCAGAGGCATAC 3’, probe 56. The PCR conditions were 50°C for 2 minutes, 95°C for 10 minutes, 40 x (95°C for 10 seconds, 60°C for 20 seconds).

### Electrophysiological analyses

For electrophysiological analyses, animals at P20 were anesthetized using 2,2,2-tribromoethanol (Sigma-Aldrich T48402) and maintained under a heat lamp throughout the procedure. Nerve conduction velocity (NCV) and compound muscle action potential amplitude were measured from sciatic nerves using a TECA Synergy N2 electromyography stimulator and subdermal steel monopolar needle electrodes. The recording electrode was positioned in the muscle in the middle of the paw, a reference electrode was positioned in between two digits, and a ground lead was inserted into the cervical musculature. Stimulation was performed using a pair of electrodes positioned at three different points: at the level of the ankle, the sciatic notch, and in the paraspinal region at the level of the iliac crest. Distal amplitude was measured from the first position (ankle) and the average from both sciatic nerves taken for each animal (unless measurements were only able to be acquired from one side, and then this was reported). NCV was calculated based on the measured distances between stimulation sites from both sciatic nerves and the average for each animal is reported (unless measurements were only able to be acquired from one side, and then this is reported).

### Statistical analyses

No power analyses were performed for this study, but our sample sizes are similar to those generally used in the field. All data are presented as the mean ± the standard error of the mean (SEM). Statistical analyses were performed using GraphPam Prism 10.0.3 and values of P < 0.05 were considered significant. The statistical tests performed for each data set are reported in the figure legends.

## Results

### Schwann cell PHB2 is essential for proper peripheral nerve development and myelination from the earliest stages

Our group previously showed that loss of Schwann cell PHB2 in mice (PHB2-SCKO) results in a radial sorting and myelination defect in the sciatic nerve, shown at post-natal days 20 and 40 (P20 and P40) (Poitelon et al., 2015). Examination of the sciatic nerve of PHB2-SCKO animals before P20 reveals that substantial developmental defects are evident as early as P5 (Fig. 1A). These animals display very limited progression through the typical peripheral nerve developmental stages as the mice age to P10 (Fig. 1B) and P20 (Fig. 1C), displaying a severe reduction in the number of myelinated axons (Fig. 1D) and by P20 numerous bundles of unsorted axons remain (Fig. 1E).

**Figure 1.**
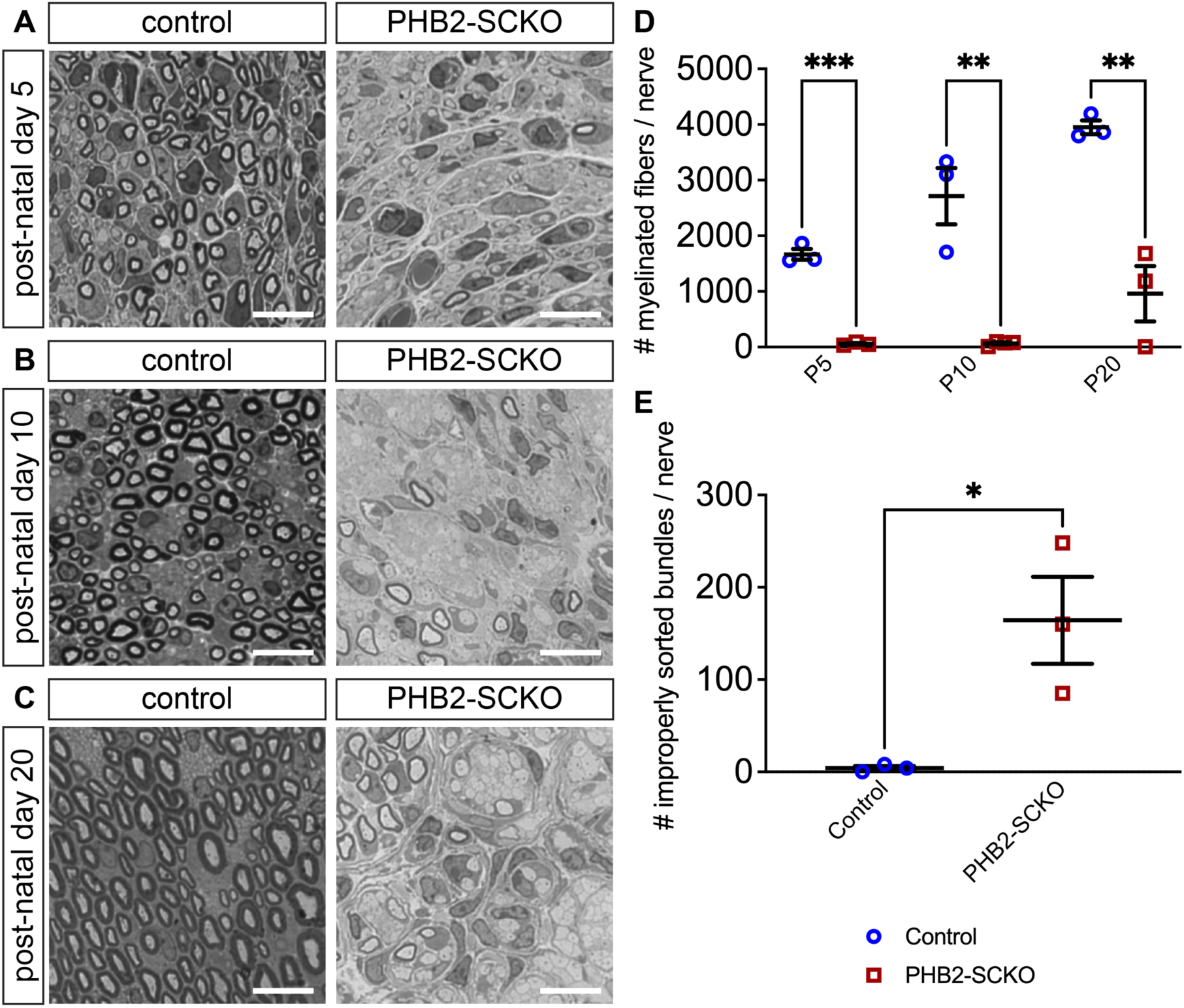
PHB2-SCKO nerves display developmental defects as early as post-natal day (P)5. Representative cross sections from the sciatic nerves of mice at P5 (A), P10 (B) and P20 (C). In controls, dark staining of the myelin sheath rings around axons is evident at all ages. In PHB2-SCKO animals there are far fewer myelinated axons and at P20 many large sized axons remain in unsorted bundles. Quantification of the number of myelinated axons per nerve cross section are shown in (D) and the number of improperly sorted bundles per nerve cross section in P20 mice in (E). *N* = 3-4 animals per genotype and each data point represents an individual animal. Data are presented as mean ± SEM and statistical significance, as calculated by unpaired t-test and in (D) corrected for multiple comparisons using the Holm-Šídák method, indicated as follows: * P < 0.05, ** P ≤ 0.01, *** P ≤ 0.001. Scale bar = 10 μm.

As PHBs play roles in multiple cell pathways (i.e. including in ERK, AKT, NF-κB and STAT3 signaling and in mitophagy) and in several sub-cellular compartments (including mitochondria, plasma membrane and nucleus) (Wang et al., 2020), we aimed to narrow down which of the PHBs’ functions contribute most to peripheral myelination. We tested several PHB modulating compounds with some specific mechanisms of action in an *in vitro* co-culture myelinating system of primary rat dorsal root ganglia (DRG) neurons and Schwann cells. We used a semi-automated system for analysis of the number of myelin internodes which we found was able to produce results in line with a manual analysis approach (Supp. Fig. 1A-C). We tested flavagline (FL3), sulfoamidine (SA1M), two melanogenin derivatives (Mel6 and Mel41), and fluorizoline (Fz), which all directly interact with PHBs to modulate their various functions, and a hydantoin derivative, an indirect agonist of PHB cytosolic functions (see Table 2 for details). While treatment with low concentrations of PHB-modulating ligands had either no effect on myelination, or, on occasion, caused a modest increase in the number of myelin internodes (Supp. Fig. 1D-O), treatment with the highest concentration of five out of six of the compounds (all but Mel6) resulted in a significant reduction in the number of myelin segments (FL3, -43% ± 10%, SA1M, -92% ± 15%, Fz, -78% ± 9%, Mel41 -58% ± 10%, HD1, -75% ± 8%, Fig. 2). Overall, while these experiments did not clearly reveal which PHB subcellular activity/ies are the most important for myelination, they do serve as pharmacological evidence supporting our *in vivo* data showing a key role for PHBs in Schwann cells.

**Figure 2.**
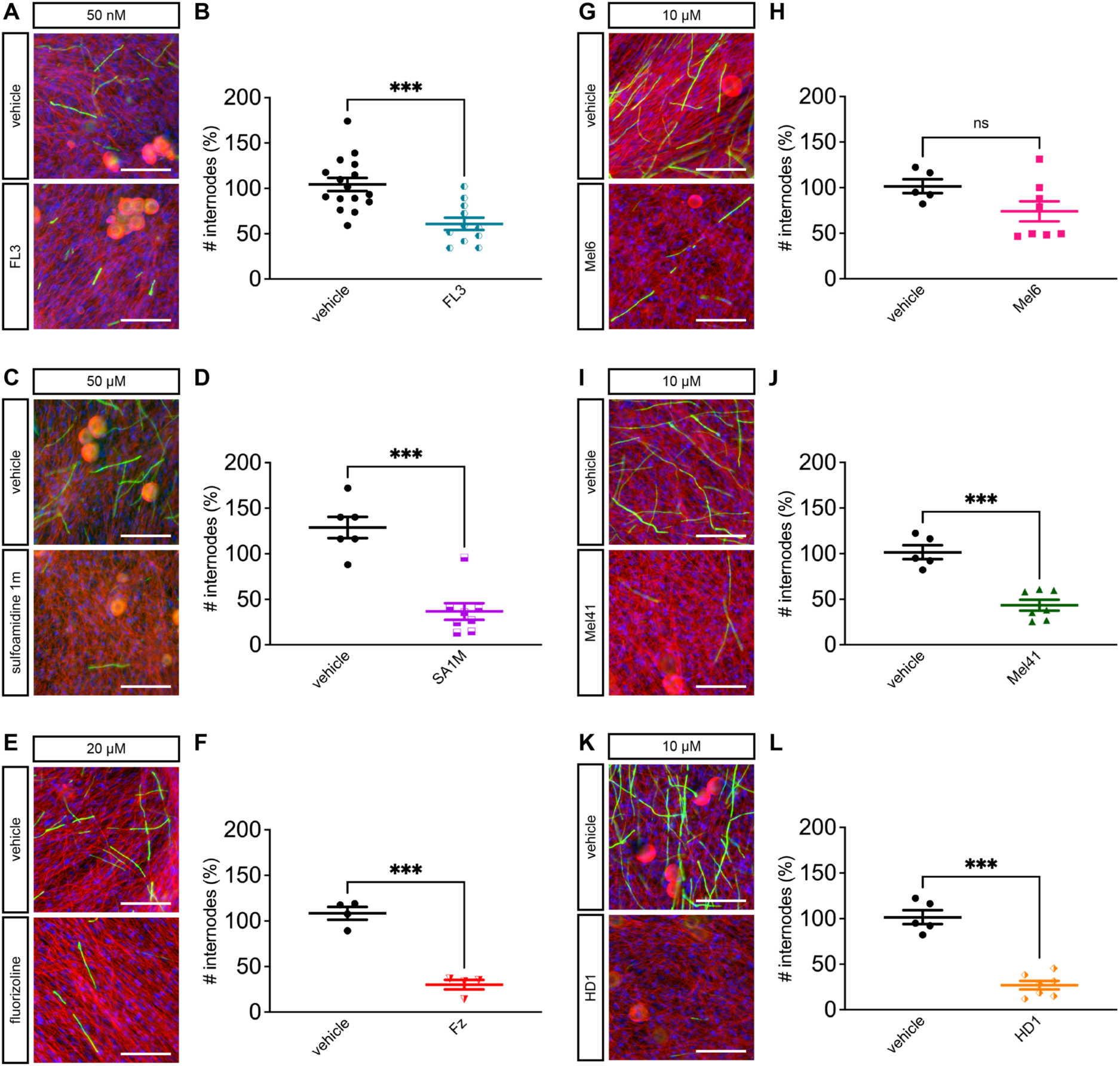
PHB-modulating compounds interfere with *in vitro* myelination. Representative immunofluorescent images and corresponding quantifications of the percentage of myelin internodes as a percentage relative to untreated controls following treatment with six different PHB-modulating compounds, at the concentrations indicated. β-III tubulin immunostaining is shown in red, myelin basic protein (MBP) in green and DAPI is in blue. Scale bar = 100 μm. Each data point represents an individual coverslip, from which the number of internodes was averaged from 3-6 random fields of view. Data were pooled from at least two independent rat preparations of neurons, other than in the fluorizoline (Fz) condition, in which all coverslips were from one preparation. Data are presented as mean ± SEM and statistical significance as calculated by unpaired t-test indicated as follows: * P < 0.05, ** P ≤ 0.01, *** P ≤ 0.001.

**Table 2:**
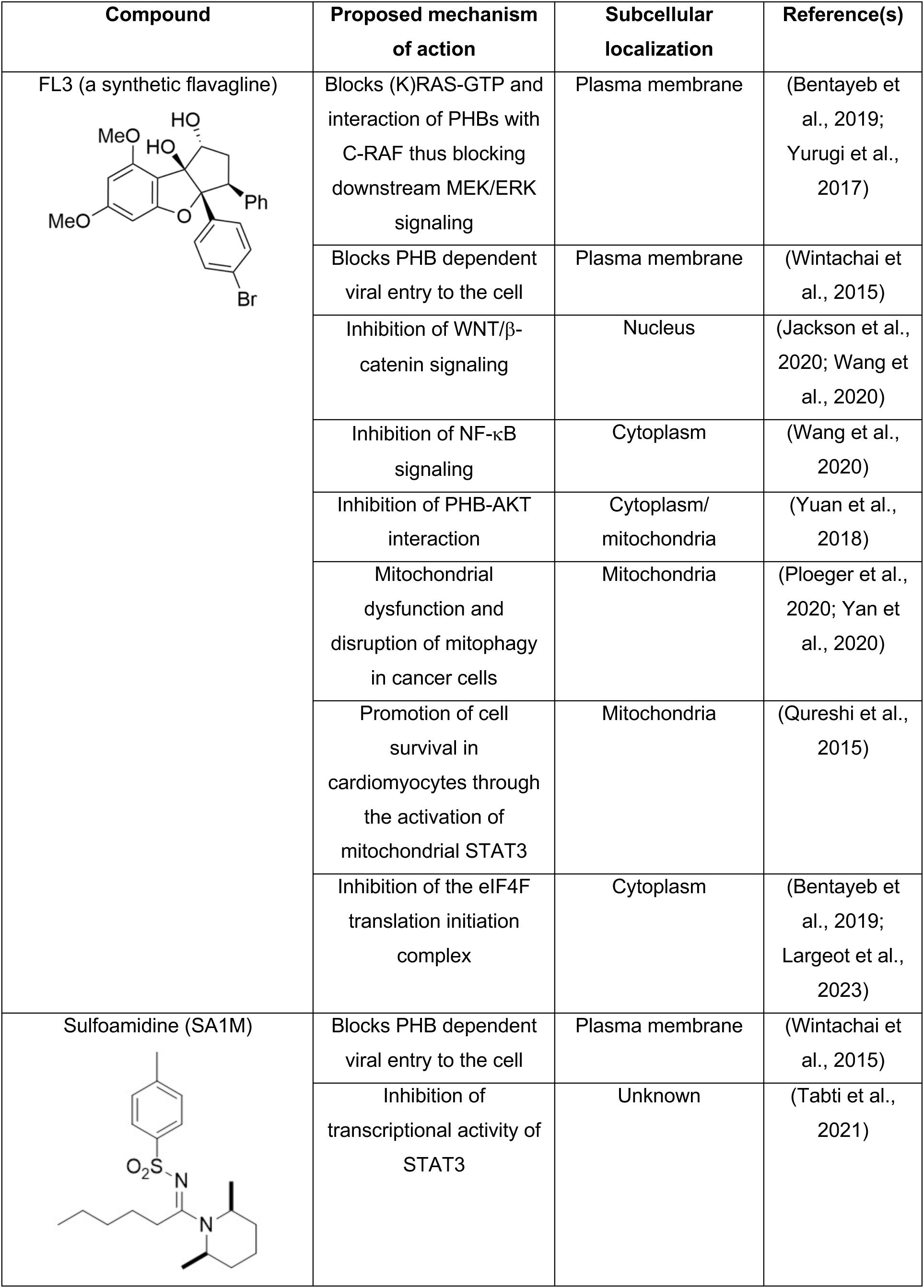
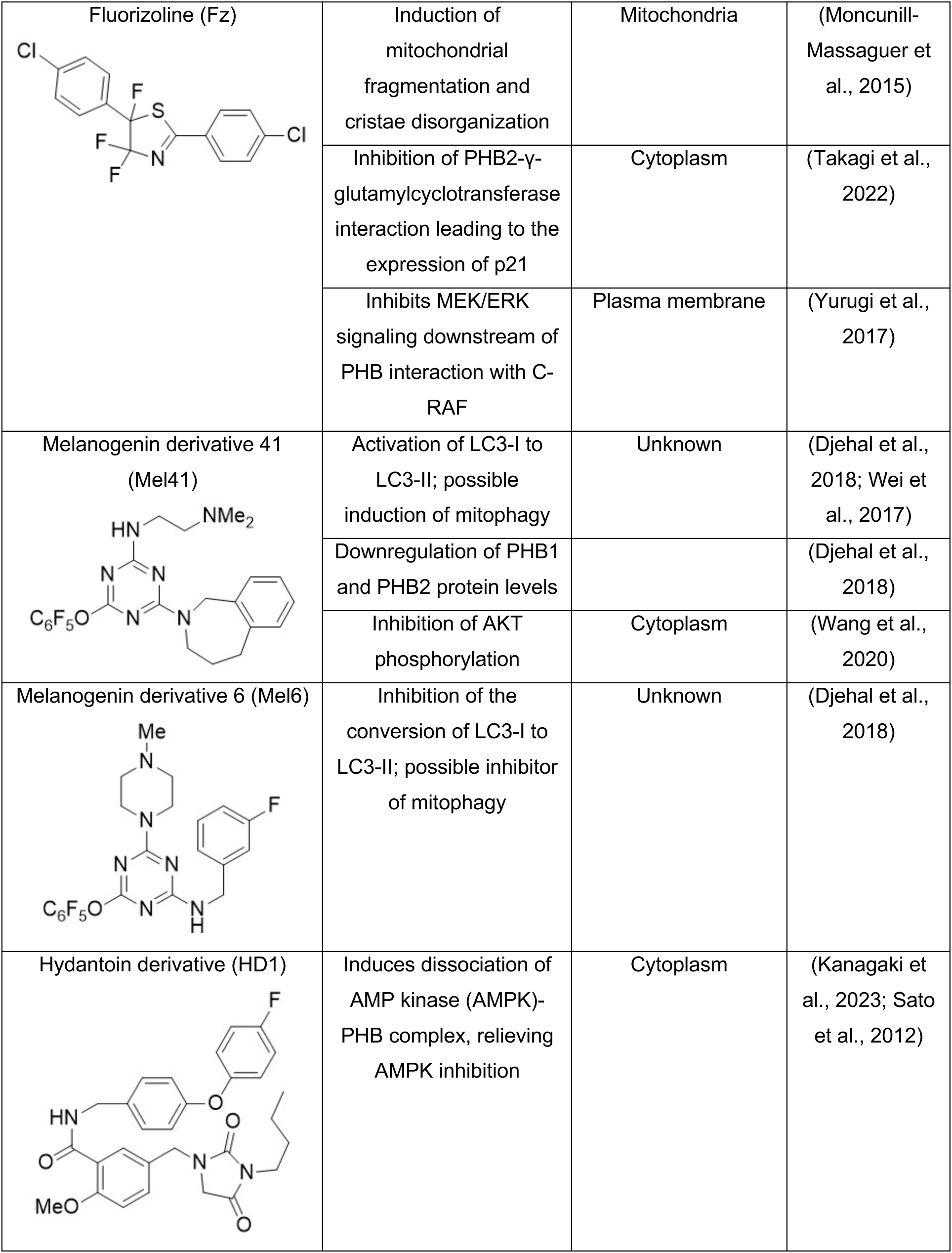
A summary of the PHB-modulating ligands tested for an effect on myelination, some of their proposed mechanisms of action relating to the PHBs, the subcellular localization of the PHBs on which they act, and the relevant references. For an in-depth review of PHB modulating ligands see (Wang et al., 2020).

| Compound | Proposed mechanism of action | Subcellular localization | Reference(s) |
| --- | --- | --- | --- |
| FL3 (a synthetic flavagline) | Blocks (K)RAS-GTP and interaction of PHBs with C-RAF thus blocking downstream MEK/ERK signaling | Plasma membrane | (Bentayeb et al., 2019; Yurugi et al., 2017) |
|  | Blocks PHB dependent viral entry to the cell | Plasma membrane | (Wintachai et al., 2015) |
| | Inhibition of WNT/β-catenin signaling | Nucleus | (Jackson et al., 2020; Wang et al., 2020) |
| | Inhibition of NF-κB signaling | Cytoplasm | (Wang et al., 2020) |
|  | Inhibition of PHB-AKT interaction | Cytoplasm/mitochondria | (Yuan et al., 2018) |
|  | Mitochondrial dysfunction and disruption of mitophagy in cancer cells | Mitochondria | (Ploeger et al., 2020; Yan et al., 2020) |
|  | Promotion of cell survival in cardiomyocytes through the activation of mitochondrial STAT3 | Mitochondria | (Qureshi et al., 2015) |
|  | Inhibition of the eIF4F translation initiation complex | Cytoplasm | (Bentayeb et al., 2019; Largeot et al., 2023) |
| Sulfoamidine (SA1M) | Blocks PHB dependent viral entry to the cell | Plasma membrane | (Wintachai et al., 2015) |
|  | Inhibition of transcriptional activity of STAT3 | Unknown | (Tabti et al., 2021) |
| Fluorizoline (Fz) | Induction of mitochondrial fragmentation and cristae disorganization | Mitochondria | (Moncunill-Massaguer et al., 2015) |
| | Inhibition of PHB2-γ-glutamylcyclotransferase interaction leading to the expression of p21 | Cytoplasm | (Takagi et al., 2022) |
|  | Inhibits MEK/ERK signaling downstream of PHB interaction with C-RAF | Plasma membrane | (Yurugi et al., 2017) |
| Melanogenin derivative 41 (Mel41) | Activation of LC3-I to LC3-II; possible induction of mitophagy | Unknown | (Djehal et al., 2018; Wei et al., 2017) |
|  | Downregulation of PHB1 and PHB2 protein levels |  | (Djehal et al., 2018) |
|  | Inhibition of AKT phosphorylation | Cytoplasm | (Wang et al., 2020) |
| Melanogenin derivative 6 (Mel6) | Inhibition of the conversion of LC3-I to LC3-II; possible inhibitor of mitophagy | Unknown | (Djehal et al., 2018) |
| Hydantoin derivative (HD1) | Induces dissociation of AMP kinase (AMPK)-PHB complex, relieving AMPK inhibition | Cytoplasm | (Kanagaki et al., 2023; Sato et al., 2012) |

### The Schwann cell PHB2 interactome is altered in the presence of neurons

Previous work indicated that PHB2 can be found in the plasma membrane of Schwann cells and especially so in response to neuronal signals (Poitelon et al., 2015). To further understand the role of PHBs in Schwann cell-neuron communication, we next sought to evaluate whether the interactions of Schwann cell PHBs with other proteins change depending on neuronal signals. To this end, we expressed a PHB2-turboID fusion construct in primary rat Schwann cells. TurboID is a 35 kDa engineered biotin ligase that rapidly biotinylates proximal proteins. Thus, proteins interacting with PHB2-turboID within an approximately 10 nm radius are tagged with biotin (Branon et al., 2018; Kim et al., 2014). As a control, primary rat Schwann cells were transfected with an unfused turboID construct (Con-turboID). Schwann cells expressing Con-turboID or PHB2-turboID were plated alone or onto primary rat DRG neurons, in the presence or absence of biotin (Fig. 3A and Supp. Fig. 2). After 2 hours, proteins were harvested from the cultures and biotinylated proteins were purified using streptavidin-affinity purification (AP). We then used liquid chromatography-mass spectrometry (LC-MS) to identify the PHB2-turboID interactors (the biotinylated proteins) in our purified pool. The expression of each protein identified was calculated relative to the total pool of biotinylated proteins for each condition within each replicate. We identified 17 proteins that interacted with PHB2-turboID either exclusively or preferentially in the presence of neurons (Fig. 3B). We also identified 14 proteins that decreased their interactions with PHB2-turboID when Schwann cells were plated with neurons, compared to Schwann cells plated alone (Fig. 3C).

**Figure 3.**
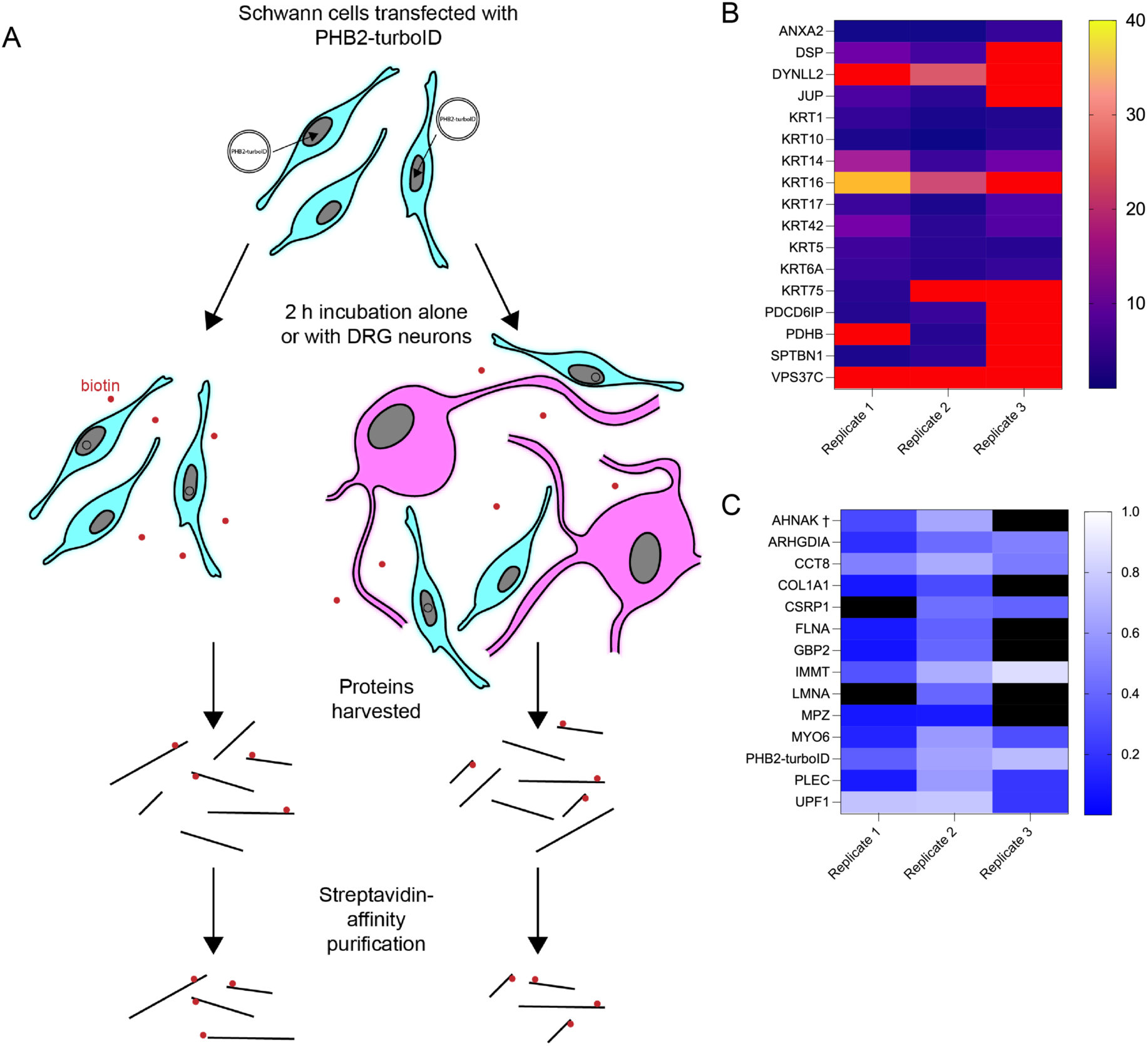
The PHB2 interactome in the Schwann cell changes depending on neuronal signals. (A) Schematic demonstrating the experiment protocol. Primary rat Schwann cells were transfected with PHB2-turboID and plated alone or with primary rat dorsal root ganglia (DRG) neurons, with or without biotin supplementation, for 2 h. Biotinylated proteins were harvested using streptavidin-affinity purification (AP) and analysed using LC-MS. (B) Proteins identified by LC-MS which interacted with PHB2-turboID to a greater extent when Schwann cells were plated with neurons, compared to when Schwann cells were plated alone. The proteins listed were enriched in their PHB2-turboID interactions in three independent replicates, and the relative total ion chromatograph (TIC) count intensity compared to when Schwann cells were plated alone is indicated by colour. Red panels indicate a protein that did not interact at all with PHB2-turboID in the Schwann cell alone condition, but was identified as a PHB2-turboID interactor when neurons were present. (C) Proteins identified by LC-MS which decreased interactions with PHB2-turboID when Schwann cells were plated with neurons, compared to when Schwann cells were plated alone. The proteins listed decreased their PHB2-turboID interaction in three independent replicates, with the relative TIC count intensity of each protein in the Schwann cells plated with neurons condition compared to Schwann cells alone, indicated by the colour of the box. Candidates who interacted with PHB2-turboID only if neurons were absent are indicated by black panels. ^†^ AHNAK fragment.

Interestingly, many of the proteins identified (Fig. 3) appear to be frequently associated with the cytoskeleton and/or scaffolding between the cytoskeleton and the plasma membrane. For example, two proteins that increased interaction with PHB2-turboID when Schwann cells were plated with neurons were spectrin β chain (SPTBN1), a subunit of the spectrin scaffolding protein which can link the actin cytoskeleton with the plasma membrane and is required for Schwann cell myelination (Goodman et al., 2019; Susuki et al., 2011) and annexin A2 (ANXA2), a protein whose expression has been previously reported to be upregulated in Schwann cell pseudopods ahead of cytoskeletal remodeling and cell migration after injury (Negro et al., 2018). In contrast, filamin A (FLNA), which can serve as an actin-binding scaffold protein, decreased interaction with PHB2-turboID when neuronal signals were present. Dynein light chain 2 (DYNLL2), a component of the dynein motor protein complex, is another cytoskeletal-associated protein that interacted with PHB2-turboID to a greater extent when Schwann cells were seeded with neurons compared to when Schwann cells were plated alone, whereas another motor protein subunit, myosin VI (MYO6) interacted less with PHB2-turboID when in the presence of neuronal signals.

Proteins identified with non-cytoskeletal functions include programmed cell death 6-interacting protein (PDCD6IP, also called ALIX or AIP1) and VPS37C, which both play a role in the endosomal sorting complexes required for transport (ESCRT) pathway (Williams & Urbe, 2007) and were found to interact with PHB2-turboID more readily, or sometimes exclusively, when neurons were present. Perhaps unsurprisingly, given the prominent role for PHBs in mitochondria, some mitochondrial proteins were identified as interacting with PHB2-turboID: pyruvate dehydrogenase E1 component subunit beta (PDHB) interacted more with PHB2-turboID in the presence of neuronal signals, whereas MICOS complex subunit MIC60 (IMMT) interacted less in the presence of neuronal signals. The key Schwann cell myelin protein myelin protein zero (MPZ) was also found to decrease interaction with PHB2-turboID in the presence of neuronal signals.

Finally, we identify four proteins associated with cell-cell contact by way of the formation of desmosomes: desmoplakin (DSP) and plakoglobin (JUP), which increased their interaction with PHB2-turboID, and AHNAK1 (also known as desmoyokin) and plectin (PLEC), which decreased interaction. A role for AHNAK1 in peripheral nerve development has been suggested previously (Salim et al., 2009) and Schwann cell plectin is thought to contribute to myelin stability (Walko et al., 2013). Notably, numerous keratin proteins were purported to increase their interaction with PHB2-turboID when Schwann cells were seeded with neurons. While keratins are frequently described as common contaminants in proteomic studies, since we detected many keratins even after filtering to remove non-specific candidates, we chose to leave them in the data set depicted in Fig. 3. A full list of LC-MS identified proteins in each experimental condition can be seen in Supp. data 1 (Excel file available from corresponding author).

### PHB2-SCKO animals may respond to radial sorting defects by modulating the AKT/mTORC1 signaling axis

Mice lacking Schwann cell PHB2 are unable to properly sort axons into one-to-one relationships with Schwann cells and subsequently cannot myelinate those axons during peripheral nerve development (Poitelon et al., 2015 and Fig. 1). Typically during peripheral nerve development, axonal NRG1-III binds to Schwann cell ERBB2/3 receptors to activate downstream signaling cascades including MAPK/ERK and PI3K/AKT/mTOR in the Schwann cell, which ultimately promote the production of transcription factors necessary for Schwann cell proliferation, maturation and myelination (Monk et al., 2015). Thus, we next sought to understand if these signaling pathways are perturbed in PHB2-SCKO animals. Using western blot, we examined protein levels in the sciatic nerves of animals aged P1, P5, P10 and P20 (Fig. 4). When activated, the MAPK signaling pathway converges on the phosphorylation of ERK. We found that ERK1/2 phosphorylation was unchanged in the sciatic nerves of PHB2-SCKO mice when compared to controls, suggesting this signaling axis does not play a role in myelination downstream of PHBs (Supp. Fig. 3A-B, Fig. 4A). Careful temporal control of PI3K-AKT-mTORC1 signaling is also required for proper peripheral myelination. High mTORC1 signaling early in development promotes immature Schwann cell proliferation via suppression of EGR2 via S6 kinase. mTORC1 signaling must subsequently be dampened for Schwann cells to terminally differentiate and trigger the onset of myelination. However, following the establishment of mature myelinating Schwann cells, mTORC1 signaling then drives myelination (Figlia et al., 2017). We assessed the phosphorylation of AKT (p-AKT) and detected an increased in p-AKT relative to total AKT levels in PHB2-SCKO nerves specifically at P5 (Supp. Fig. 3C-D, Fig. 4B). Next, we measured mTORC1 signaling by examining total and phosphorylated levels of molecules targeted by mTORC1, namely, the S6 ribosomal protein (S6RP) and 4E-BP1. At P5 we see evidence of an increase in mTORC1 signaling in the sciatic nerve of PHB2-SCKO animals, compared to control mice, as demonstrated by an increase in the phosphorylation of the S6RP and 4EBP1, relative to total protein levels (Supp. Fig. 3E-H, Fig. 4C-D). Interestingly, at P20 we observe an increase in total S6RP, total 4E-BP1 and p-4E-BP1 levels (Supp. Fig. 3E, G, H).

**Figure 4.**
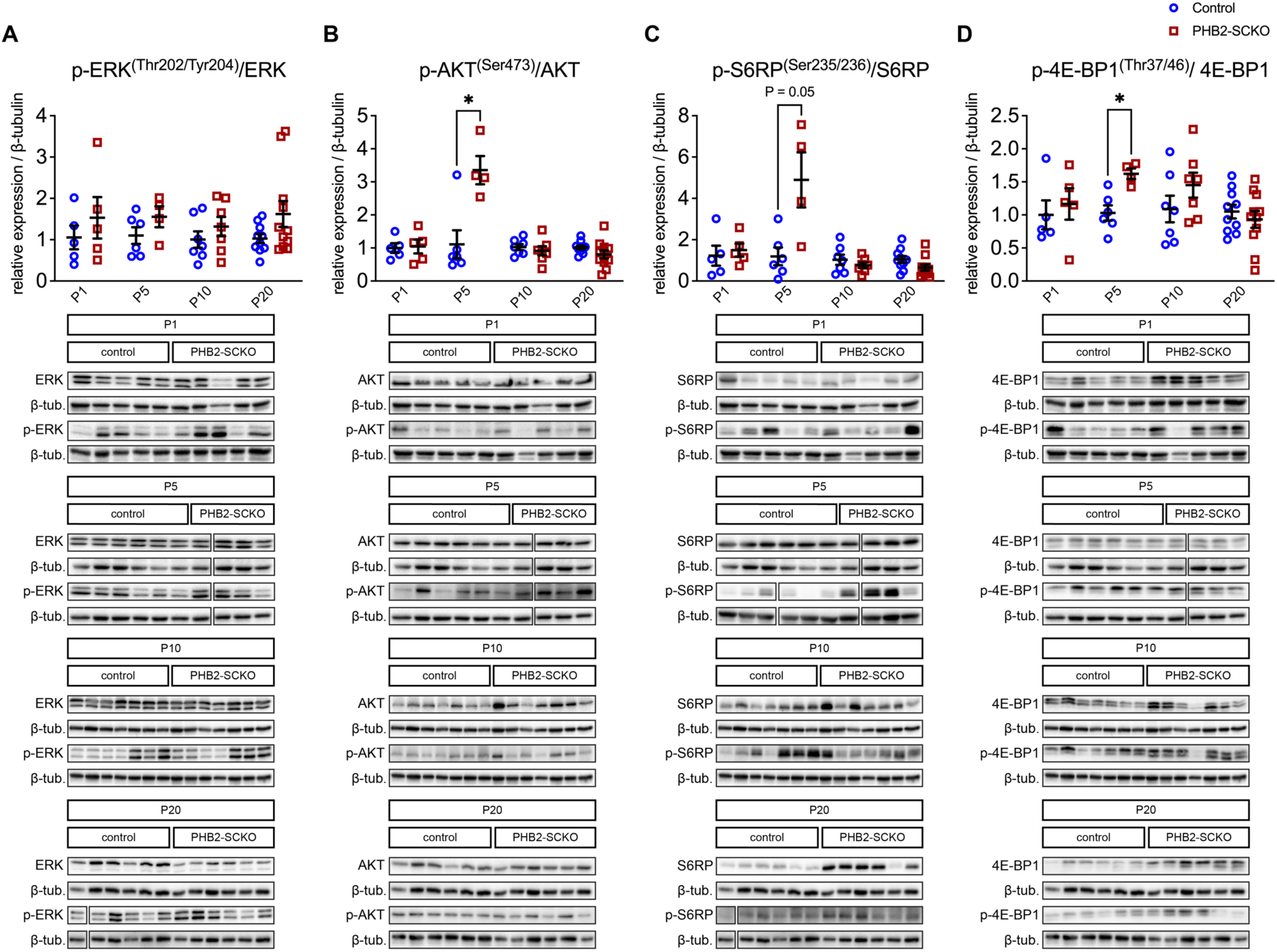
AKT and mTORC1 signaling is elevated at P5 in PHB2-SCKO nerves. (A) Quantification (upper panel) of western blots (lower panels) probing for total ERK, phosphorylated ERK (p-ERK) and β-tubulin (loading control) in sciatic nerve lysate from P1, P5, P10 and P20 control and PHB2-SCKO animals. (B) Quantification (upper panel) of western blots (lower panels) probing for total AKT, phosphorylated AKT (p-AKT) and β-tubulin (loading control) in sciatic nerve lysate from P1, P5, P10 and P20 control and PHB2-SCKO animals. (C) Quantification (upper panel) of western blots (lower panels) probing for total S6RP, phosphorylated S6RP (p-S6RP) and β-tubulin (loading control) in sciatic nerve lysate from P1, P5, P10 and P20 control and PHB2-SCKO animals. (D) Quantification (upper panel) of western blots (lower panels) probing for total 4E-BP1, phosphorylated 4E-BP1 (p-4E-BP1) and β-tubulin (loading control) in sciatic nerve lysate from P1, P5, P10 and P20 control and PHB2-SCKO animals. Each data point represents an individual animal, except for in the P1 and P5 peripheral nerve lysates, when multiple nerves from animals with the same genotype were pooled, and each data point represents an independent pool of nerves. Expression of the protein of interest was first calculated relative to the loading control β-tubulin and then phosphorylated relative to total levels, as total and phosphorylated protein levels were probed for on different membranes. Expression was then normalised to the average relative expression of the controls, for each age. In some cases the same membrane was probed sequentially for multiple signaling proteins. Uncropped blots are shown in Supp. Fig. 7. Data are presented as mean ± SEM and statistical significance, calculated by unpaired t-test and corrected for multiple comparisons using the Holm-Šídák method, indicated as follows: * P < 0.05, ** P ≤ 0.01, *** P ≤ 0.001.

In summary, we did not detect widespread changes in regulation of either the MAPK/ERK or PI3K/AKT/mTORC1 signaling cascades, suggesting that changes in these signaling molecules are unlikely to underlie the severe and early onset developmental defects present in PHB2-SCKO animals. We hypothesize that the narrow window of change at P5 in the AKT/mTORC1 signaling axis reflects a response to the radial sorting defect in these nerves, as opposed to a casual relationship. We propose that Schwann cells lacking PHB2 respond to signals indicating that radial sorting is incomplete by increasing p-AKT and mTORC1 signaling to suppress EGR2 and delay the onset of myelination. To test this hypothesis, we next examined the levels of the transcription factor EGR2, and other transcription factors, in our conditional knockout nerves.

### Loss of PHB2 in Schwann cells alters the expression of myelin transcriptional regulators

The PI3K-AKT-mTORC1 signaling axis has been linked to control of the transcription factor EGR2 (also known as KROX20, a master regulator of peripheral myelination) (Figlia et al., 2017). Since we saw some changes in AKT and mTORC1 signaling in the peripheral nerve of PHB2-SCKO animals (Fig. 4), we also tested for changes in the transcription factors governing Schwann cell development and myelination. We found that PHB2-SCKO animals have significantly less EGR2 protein in their sciatic nerves at P5 and P10, when compared to controls of the same age (Fig. 5A-B). This was also evident by immunohistochemistry of the nerve at P5, where PHB2-SCKO animals have 20% ± 5% fewer EGR2 positive nuclei in their sciatic nerves than controls (Fig. 5C-D). The sciatic nerve of PHB2-SCKO animals is also persistently thinner and visibly more transparent than in control animals. POU3F1 (also known as OCT6), another important Schwann cell transcription factor, governs EGR2 expression (Blanchard et al., 1996; Ghislain et al., 2002). In PHB2-SCKO animals we also detect depletion of POU3F1, in this case starting as early as P1 in the sciatic nerve, and evident up to P10 (Fig. 5E-F). In line with this finding, immunohistochemistry of the sciatic nerve at P5 demonstrated a reduction of 31% ± 7% in the number of POU3F1 positive cells in PHB2-SCKO nerves compared to controls (Fig. 5G-H). Interestingly, despite being reduced in PHB2-SCKO nerves when compared to controls from P1 to P10, at P20, PHB2-SCKO sciatic nerves have an increase in the amount of POU3F1 protein comparatively to controls (Fig. 5E-F). Similarly, the transcription factor POU3F2 (also known as BRN2), which is functionally similar to POU3F1 (Jaegle et al., 2003), follows a near identical expression pattern to POU3F1, whereby it is reduced in PHB2-SCKO sciatic nerves when compared to controls at P1 through to P10, but increases comparatively to control nerves at P20 (Fig. 5I-J). To summarize, from P1 to P10, the protein levels of the transcription factors POU3F1 and POU3F2 are reduced in PHB2-SCKO nerves compared to their control counterparts, and subsequently, from P5 to P10, the protein levels of EGR2 are also reduced in PHB2-SCKO nerves, yet at P20 the protein levels of all three transcription factors are elevated in the conditional knockout nerves.

**Figure 5.**
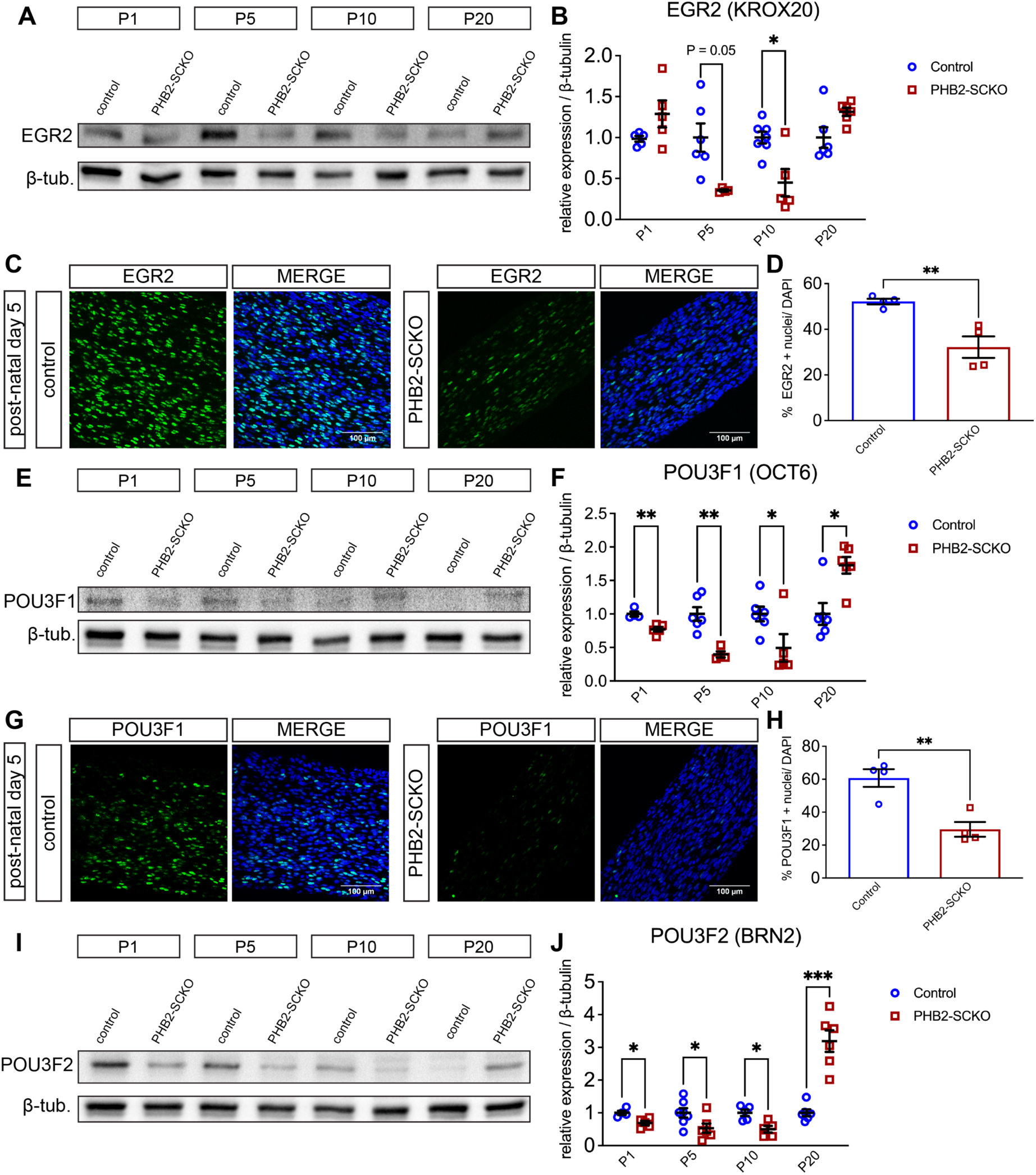
Key Schwann cell transcription factors are dysregulated in PHB2-SCKO nerves. (A) Representative western blot showing EGR2 and β-tubulin (loading control) levels in peripheral nerve lysate from P1, P5, P10 and P20 control and PHB2-SCKO animals. (B) Quantification of EGR2 levels from western blots. (C) Representative images showing immunohistochemistry of EGR2 (green) in P5 control and PHB2-SCKO longitudinal sciatic nerve sections. DAPI is shown in blue. (D) Quantification of the number of EGR2 positive nuclei (as a percentage of total nuclei) shown in (C). (E) Representative western blot showing POU3F1 (OCT6) and β-tubulin (loading control) levels in peripheral nerve lysate from P1, P5, P10 and P20 control and PHB2-SCKO animals. (F) Quantification of the POU3F1 levels from western blots as per (E). (G) Representative images showing immunohistochemistry of POU3F1 (green) in P5 control and PHB2-SCKO sciatic nerve longitudinal sections. DAPI is shown in blue. (H) Quantification of the number of POU3F1 positive nuclei (as a percentage of total nuclei) shown in (G). (I) Representative western blot showing POU3F2 (BRN2) and β-tubulin (loading control) levels in peripheral nerve lysate from P1, P5, P10 and P20 control and PHB2-SCKO animals. (J) Quantification of the POU3F2 levels from western blots as per (I). Each data point represents an individual animal, except for in the P1 and P5 peripheral nerve lysates, when multiple nerves from animals with the same genotype were pooled, and each data point represents an independent pool of nerves. Expression of the protein of interest was calculated relative to the loading control β-tubulin then normalised to the average relative expression of the controls, for each age. Data are presented as mean ± SEM. In (B), (F) and (J) statistical significance was calculated by unpaired t-test and corrected for multiple comparisons using the Holm-Šídák method. In (D) and (H) statistical significance was calculated by unpaired t-test. Statistical significance is represented on the graphs as follows: * P < 0.05, ** P ≤ 0.01, *** P ≤ 0.001.

A possible explanation for the reduction in the overall levels of pro-myelinating transcription factors seen in Fig. 5, is that there are fewer Schwann cells in PHB2-SCKO peripheral nerves relative to their control counterparts. SOX10 is a transcription factor expressed by all cells derived from the neural crest, and it is necessary for peripheral glia specification (Britsch et al., 2001). PHB2-SCKO sciatic nerves display lower SOX10 levels than controls at P10, and reduced nominal levels at P1, P5, and P20, although statistical significance was not reached, as measured by western blot (Fig. 6A-B). There is also 18% ± 6% fewer SOX10 positive nuclei in the sciatic nerve of PHB2-SCKO animals compared to controls at P5, as measured by immunohistochemistry (Fig. 6C-D). When examining the ability of PHB2-null Schwann cells to proliferate using the marker of proliferation MKI67, we found that only 6% ± 2% of SOX10 positive cells in the sciatic nerve of P5 PHB2-SCKO animals also express MKI67, compared to 33% ± 1% in controls (Fig. 6C and E), suggesting that PHB2-null Schwann cells do not proliferate in the same manner as Schwann cells in the wildtype nerve.

**Figure 6.**
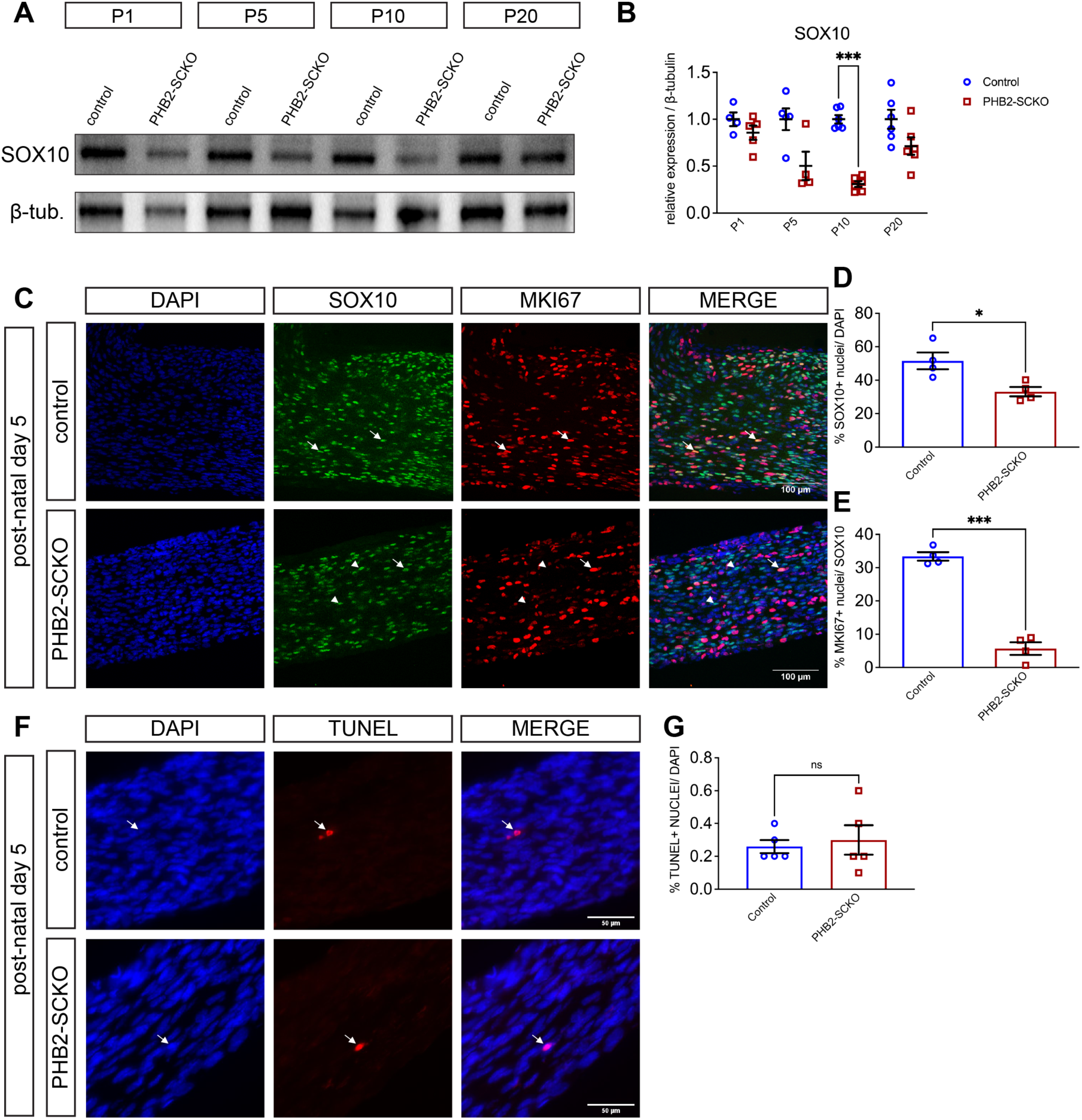
Schwann cells in PHB2-SCKO nerves do not proliferate effectively. (A) Representative western blot showing SOX10 and β-tubulin (loading control) levels in peripheral nerve lysate from P1, P5, P10 and P20 control and PHB2-SCKO animals. (B) Quantification of SOX10 levels from western blots. Expression of SOX10 was calculated relative to the loading control β-tubulin then normalised to the average relative expression of the controls, for each age. (C) Representative images showing immunohistochemistry of SOX10 (green) and proliferation marker MKI67 (red) in P5 control and PHB2-SCKO sciatic nerve longitudinal sections. DAPI is shown in blue. Arrows indicate proliferating Schwann cells (SOX10 and MKI67 positive). Arrow heads indicate Schwann cells not proliferating (SOX10 positive, MKI67 negative). Scale bar = 100 μm. (D) Quantification of the number of SOX10 positive nuclei (as a percentage of total nuclei) shown in (C). (E) Quantification of the number of proliferating Schwann cells, calculated as the number of MKI67 positive nuclei as a percentage of SOX10 positive nuclei, as shown in (C). (F) Representative images showing TUNEL staining (red) in P5 control and PHB2-SCKO sciatic nerve longitudinal sections. DAPI is shown in blue. Arrows indicate TUNEL positive cells dying by apoptosis. Scale bar = 50 μm. (G) Quantification of the number of TUNEL positive nuclei (as a percentage of total DAPI positive nuclei). Each data point represents an individual animal, except for in the P1 and P5 peripheral nerve lysates, when multiple nerves from animals with the same genotype were pooled, and each data point represents an independent pool of nerves. Data are presented as mean ± SEM. In (B) statistical significance was calculated by unpaired t-test and corrected for multiple comparisons using the Holm-Šídák method. In (D), (E) and (G) statistical significance was calculated by unpaired t-test. Statistical significance is represented on the graphs as follows: * P < 0.05, ** P ≤ 0.01, *** P ≤ 0.001.

The overall Schwann cell number could also be affected by Schwann cell death. However, TUNEL staining, which marks apoptotic nuclei, in P5 sciatic nerves shows no differences in apoptosis between PHB2-SCKO animals and controls (Fig. 6F-G). This suggests that Schwann cell death via apoptosis is not contributing to the reduction in Schwann cell number in the sciatic nerve of PHB2-SCKO animals at P5 (Fig. 6A-D).

The transcription factor JUN (also known as c-JUN) is thought to be a negative regulator of myelination, most important for the peripheral nerve regeneration process, where it instructs Schwann cells to trans-differentiate from myelinating or Remak Schwann cells into repair Schwann cells (Arthur-Farraj et al., 2012; Parkinson et al., 2008). In PHB2-SCKO sciatic nerves a persistent and approximately two-to three-fold elevation in JUN protein levels is detected from P5 to P20 compared to controls (Fig. 7A-B). An increase of 22% ± 15% (P = 0.19) of JUN-positive nuclei in the sciatic nerve is also seen by immunohistochemistry at P5 (Fig. 7C-D).

**Figure 7.**
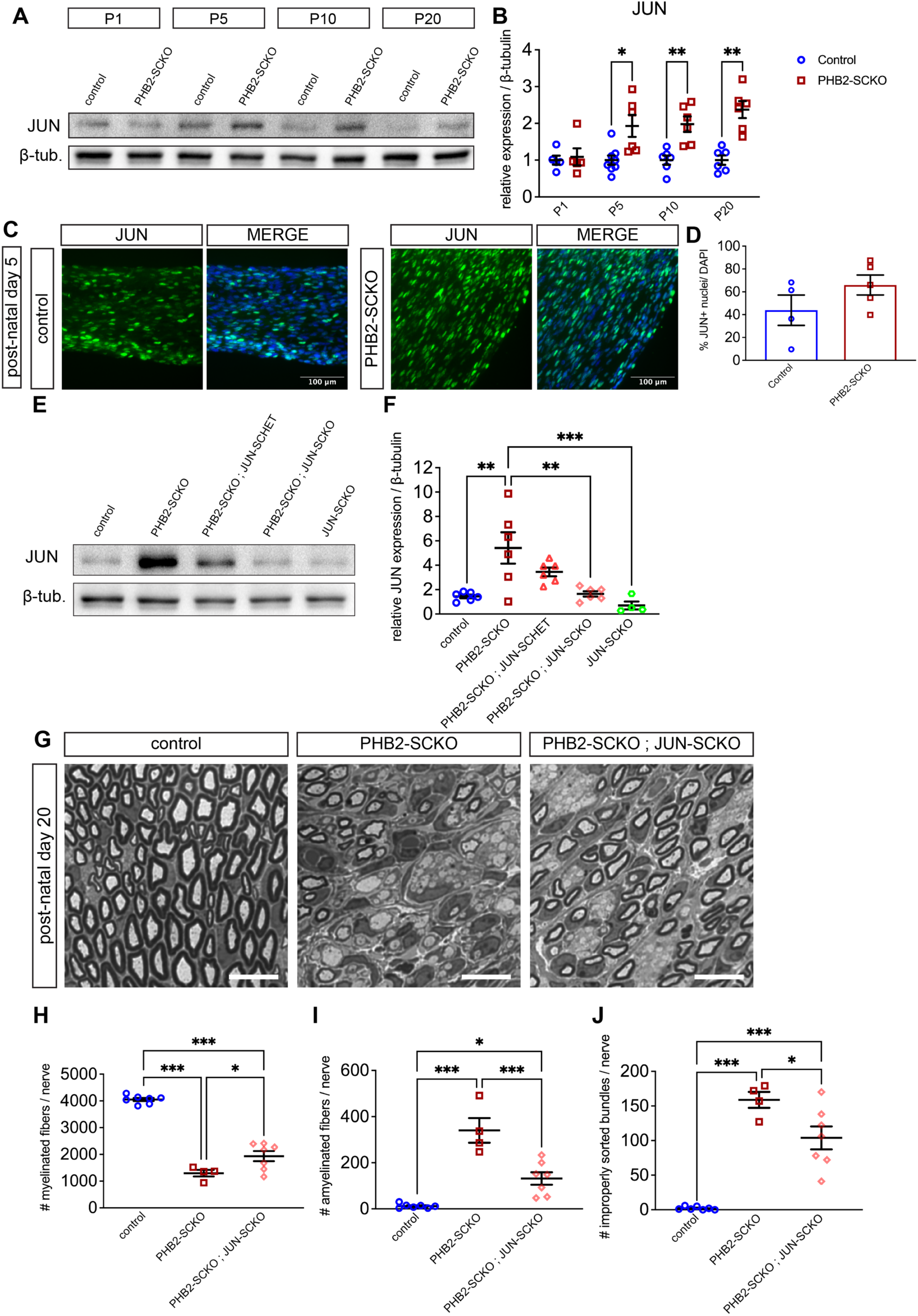
JUN-SCKO confers a partial rescue of the developmental defect in PHB2-SCKO animals. (A) Representative western blot showing JUN and β-tubulin (loading control) levels in peripheral nerve lysate from P1, P5, P10 and P20 control and PHB2-SCKO animals. (B) Quantification of the JUN levels from western blots as per (A). Expression of JUN was calculated relative to the loading control β-tubulin then normalised to the average relative expression of the controls, for each age. (C) Representative images showing immunohistochemistry of JUN (green) in P5 control and PHB2-SCKO sciatic nerve longitudinal sections. DAPI is shown in blue. (D) Quantification of the number of JUN positive nuclei (as a percentage of total nuclei) shown in (C). (E) Representative western blot showing JUN and β-tubulin (loading control) levels in peripheral nerve lysate from P20 control, PHB2-SCKO, PHB2-SCKO ; JUN-SCHET, PHB2-SCKO ; JUN-SCKO and JUN-SCKO animals. (F) Quantification of JUN levels from western blots. Expression of JUN was calculated relative to the loading control β-tubulin then normalised to the average relative expression of a control reference that was run on every blot. (G) Representative cross sections from P20 sciatic nerves. Scale bar = 10 μm. (H-J) Quantification of the number of myelinated fibers (H), amyelinated fibers (I) and improperly sorted bundles (J) from P20 sciatic nerve cross sections. Each data point represents an individual animal, except for in the P1 and P5 peripheral nerve lysates, when multiple nerves from animals with the same genotype were pooled, and each data point represents an independent pool of nerves. Data are presented as mean ± SEM. In (B) statistical significance was calculated by unpaired t-test and corrected for multiple comparisons using the Holm-Šídák method. In (D) statistical significance was calculated by unpaired t-test. In (F), (H), (I) and (J) statistical significance was calculated by one-way ANOVA: (F) F (4, 23) = 8.076, P < 0.001, (H) F (2, 15) = 100.7. P < 0.001, (I) F (2, 15) = 31.40, P < 0.001, (J) F (2, 15) = 40.52, P < 0.001 and the statistical significance indicated on the graphs was determined by Tukey’s multiple comparisons tests. * P < 0.05, ** P ≤ 0.01, *** P ≤ 0.001.

Finally, in PHB2-SCKO animals, as well as detecting the expected reduction in the mRNA levels of PHB2 in the sciatic nerve (Supp. Fig. 4A), we also observe a reduction in the mRNA levels of PHB1 (Supp. Fig. 4B). Thus, we tested to see if the transcription factors dysregulated in PHB2-SCKO animals, are also changed in the nerves of PHB1-SCKO mice. Indeed, at P5 we found a decrease in EGR2 (P = 0.11), a significant decrease in POU3F2, a significant decrease in SOX10 and a significant increase in JUN levels in PHB1-SCKO sciatic nerves compared to control counterparts (Supp. Fig. 4C, D, E and G).

### Removal of JUN from Schwann cells partially restores the developmental defect in PHB2-SCKO animals

We then investigated the causative relationship between the observed changes in transcription factors. Since JUN levels were elevated in the PHB2-SCKO animals from as early as P5 (Fig. 7A-D), we tested whether genetically lowering JUN levels in PHB2-SCKO animals can restore any of the peripheral nerve developmental defects caused by lack of Schwann cell PHB2 (Fig. 1 and (Poitelon et al., 2015)). To reduce JUN levels in Schwann cells we crossed floxed JUN mice (Behrens et al., 2002) with PHB2-SCKO mice, so that mice carrying the P0-Cre transgene express only one (JUN-SCHET) or no copies of the JUN gene (JUN-SCKO) in Schwann cells. Western blot of sciatic nerve lysate confirms that at P20, JUN levels are reduced in PHB2-SCKO ; JUN-SCHET mice, and significantly so in PHB2-SCKO ; JUN-SCKO or JUN-SCKO mice, compared to PHB2-SCKO alone (Fig. 7E-F; the low levels of JUN detected in these animals is likely contributed by peripheral nerve cell types other than Schwann cells).

Examination of the morphology of the sciatic nerve in these animals at P20 suggests that Schwann cells lacking both PHB2 and JUN can radially sort and myelinate axons more effectively than Schwann cells lacking PHB2 only (Fig. 7G). Indeed, a small increase in the number of myelinated fibers (Fig. 7H), decrease in the number of amyelinated fibers (Fig. 7I), and decrease in the number of improperly sorted bundles (Fig. 7J) suggests removal of JUN confers a partial but statistically significant rescue of the PHB2-SCKO phenotype. In line with previous findings, Schwann cell ablation of JUN alone (Arthur-Farraj et al., 2012; Della-Flora Nunes, Wilson, Hurley, et al., 2021) has no effect on the parameters of developmental myelination we examined (Supp. Fig. 5A-D). Removal of one copy of JUN in the PHB2-SCKO mice is not sufficient to rescue the radial sorting defect (Supp. Fig. 5A-D).

Investigation by electron microscopy of the sciatic nerve (Supp. Fig. 5E) reveals that the average number of axons remaining in improperly sorted bundles in PHB2-SCKO animals with genetically reduced JUN levels remains the same (Supp. Fig. 5H), as does the distribution of unsorted bundle size (Supp. Fig. 5J). Electrophysiological assessment of these animals at P20 show that while PHB2-SCKO animals display a severe reduction in nerve conduction velocity and amplitude of compound muscle action potentials, removing JUN from Schwann cells confers no statistically significant effect, either detrimental or improvement, in these animals (Supp. Fig. 5J-K). Western blot analysis of the sciatic nerves of PHB2-SCKO ; JUN-SCHET and PHB2-SCKO ; JUN-SCKO mice at P5 show that despite lowering JUN levels (Supp. Fig. 6A and F), the reduction in transcription factors EGR2, POU3F2, SOX10, and POU3F1 measured in PHB2-SCKO animals is not significantly altered (Supp. Fig. 6A-E). Thus, our results show that genetically removing JUN from Schwann cells confers a partial but statistically significant improvement of the development defect of PHB2-SCKO animals which appears to be independent of an effect on the low levels of the pro-myelinating transcription factors.

## Discussion

In this study, we build on our understanding of the PHB proteins in Schwann cell biology. We present both genetic and pharmacological evidence confirming the crucial role for PHBs in myelination. Further, we show that the PHB2 interactome changes in response to the presence of neurons, particularly with respect to cytoskeletal- and cell-cell adhesion-associated proteins. We find that Schwann cells lacking PHB2 do not readily proliferate and, finally, we report multiple transcription factor changes in our PHB2-SCKO animal model and present evidence suggesting that JUN is required for the full PHB2-SCKO phenotype.

We confirm that PHB2-SCKO mice have a severe developmental delay in their peripheral nerves, presenting as defective radial sorting and, in turn, a drastic reduction in the number of myelinated axons. In line with this, myelination is inhibited by several PHB-modulating ligands, with five out of the six tested resulting in a depletion in the formation of myelin internodes *in vitro*. Curiously, three of the compounds, FL3, SA1M, and Mel6, increase the number of myelinated internodes at some of the lower concentrations tested. Myelination is a temporally sensitive process and it is possible that dose-specific effects of some of these ligands on certain signaling pathways could cause changes in Schwann cell development and myelin wrapping that result in an increase in myelination. That being said, the broad effects of the drugs we tested make it difficult to pinpoint precisely the signaling pathway(s) involved. Nevertheless, these experiments complement what we see *in vivo*, that interfering with typical PHB activity has radical consequences on the Schwann cell’s ability to myelinate the axon.

Using a turboID-based screen, we found that the protein interactome of PHB2 changes depending on the presence of axons. Two desmosome-associated proteins, desmoplakin and plakoglobin, were found to interact more readily with PHB2-turboID when Schwann cells were plated with neurons, compared to when Schwann cells were plated alone and another two desmosome-associated proteins, AHNAK, and plectin (previously reported as a PHB interactor (Zhu et al., 2010)), were found to decrease interactions with PHB2-turboID when neurons were present. Desmosomes are not widely described in Schwann cells, although a historic study reports desmosome-like structures by electron microscopy during the formation of the myelin sheath (Gamble & Gosset, 1966). Interestingly, the polarity demonstrated by Schwann cells has been likened to epithelial cells (Arthur-Farraj et al., 2017; Bunge & Bunge, 1983) and epithelial cells are known to form desmosomes. Several proteins associated with the cytoskeleton and/or connecting the cytoskeleton with the plasma membrane were also particularly enriched in the pool of proteins that Schwann cell PHB2-turboID interacted with more or less with, in the presence of neuronal signals. This includes ANXA2 (previously reported as a PHB interactor (Bacher et al., 2002; Kolonin et al., 2004; Salameh et al., 2016)), SPTBN1, AHNAK, and FLNA. This could indicate a role for PHB2 in the colossal cytoskeletal rearrangements required for Schwann cells to sort axons during the process of radial sorting and to wrap them in layers of myelin sheath. Interestingly, FL3, one of the PHB modulating compounds we found to decrease myelination *in vitro*, has been reported to cause changes in ANXA2 levels and sub-cellular localization (Grindheim et al., 2023).

In exploring some of the key signaling molecules known to be important in Schwann cells during peripheral nerve development, we find that MEK/ERK signaling is not changed in PHB2-SCKO animals, whereas the PI3K/AKT/mTORC1 signaling pathway is dysregulated in PHB2-SCKO animals specifically at P5, as measured by the phosphorylation of AKT and downstream mTORC1 targets S6RP and 4E-BP1. While the phosphorylation of 4E-BP1 is increased relative to total 4E-BP1, this could reflect a slight decrease in total 4E-BP1 levels. Nevertheless, increased signaling of the PI3K/AKT/mTORC1 signaling axis appears to be transient and thus unlikely to be the underlying cause of the developmental defect in these animals. One hypothesis is that the elevation of mTORC1 signaling in the sciatic nerves of PHB2-SCKO animals at P5 may be a secondary effect, in response to disturbed radial sorting. As elevation in mTORC1 signaling prevents the onset of myelination, an increase in this signaling axis may act to prevent myelination of the improperly sorted nerve. Interestingly, at P20 we observe an increase in total S6RP, total 4E-BP1 and p-4E-BP1 levels. This is in line with findings in PHB1-SCKO nerves at the same age (Della-Flora Nunes, Wilson, Hurley, et al., 2021) and we suggest may be a response to the accumulation of mitochondrial damage and onset of demyelination, rather than a reflection of developmental defects.

Instead, what we do find in PHB2-SCKO nerves is a marked reduction in the number of Schwann cells (determined by SOX10, a marker of cells of the neural crest lineage) and that very few SOX10-positive cells are proliferating, at least at P5. Importantly, cell death by apoptosis does not appear to contribute to the reduction in the number of Schwann cells in the peripheral nerves of PHB2-SCKO animals. From P1, the earliest age we examined, we found a decrease in the transcription factors POU3F1 and POU3F2 in the peripheral nerves of PHB2-SCKO animals compared to controls. In turn, these transcription factors regulate the expression of EGR2, the master transcriptional regulator of many myelin genes required for peripheral myelination (including *Gjpb1*, *Mbp, Mpz* and *Pmp22* (Nagarajan et al., 2001)). In line with this, we find that from P5 EGR2 expression is also depleted in PHB2-SCKO nerves. At P20, we find the levels of these three transcription factors to instead be elevated in PHB2-SCKO nerves, compared to controls, indicating either a delay in the developmental progress of Schwann cells or a reactionary overcompensation of expression in a bid to manage the developmental block in the peripheral nerves. While mice lacking POU3F1 and POU3F2 in Schwann cells present with a delay in myelination, a radial sorting defect to the extent of in PHB2-SCKO animals is not as evident (Jaegle et al., 2003), suggesting that dysregulation of these transcription factors may not be the only contributing factor to the phenotype observed in our animals.

From P5, we also see an elevation in the transcription factor JUN, which persists through to P20. JUN is a negative regulator of myelin genes, important in the transdifferentation of Schwann cells to repair Schwann cells following nerve injury. While JUN may not contribute to a healthily developing nerve, since neither we nor others see an overt effect of conditionally ablating JUN from Schwann cells on the development of myelin (Arthur-Farraj et al., 2012; Della-Flora Nunes, Wilson, Hurley, et al., 2021), it may play a role in the context of improper development, as in our PHB2-SCKO model. It has been previously shown that forced expression of JUN suppresses EGR2 but not POU3F1 expression (Parkinson et al., 2008). Here, we show that a reduction in POU3F1 levels is evident in PHB2-SCKO nerves before EGR2 depletion. Thus, while removing JUN confers a partial but significant improvement in the number of axons that are myelinated in our model, it cannot fully restore either the cellular or molecular defects associated with removal of PHB2. Moreover, since removing PHB1 and PHB2 from Schwann cells causes demyelination (Della-Flora Nunes, Wilson, Marziali, et al., 2021 and unpublished data) it is complex to assess exactly whether the effect of JUN-SCKO on the number of myelinated axons is a result of JUN’s effect in the developing nerve or a protective effect of JUN in some axons which may have started to demyelinate already at P20, as is the case in our previous study of PHB1-SCKO animals (Della-Flora Nunes, Wilson, Hurley, et al., 2021).

In this study, we confirm the crucial role for PHBs in Schwann cells and peripheral nerve development and we shed light on the underlying mechanism. Our work reveals that PHBs are required for proper Schwann cell proliferation and differentiation. One possibility is that PHBs play a role in the cytoskeletal rearrangements that are required for Schwann cell-axon interaction, in turn permitting a cascade of signaling pathways that allow for the transcriptional changes that promote the proper formation of myelinating and Remak Schwann cells. Our data also suggests that removing JUN from Schwann cells may help to promote Schwann cell myelination in the developing nerve, at least in the context of dysregulated development.

## Acknowledgements

MLF and this work was funded by grant NIH-NINDS-R01NS100464. YP was funded by grant NIH-NINDS-R01NS110627. SM was funded by grant NIH-NINDS-F30NS125928. We are extremely grateful to Conlan Kreher for assistance with FIJI macros. EGR2 (KROX20) and POU3F1 (OCT6) antibodies were generous gifts from Dr. Dies Mejer. The MBP hybridoma was a generous gift from Dr. Judith Grinspan. We thank Dr. Bert W. O’Malley (Baylor College of Medicine) for *Phb1* floxed mice, and Dr. Erwin F Wagner (Medical University of Vienna) for the *Jun* floxed mice.

---

**Supplementary Figure 1.**
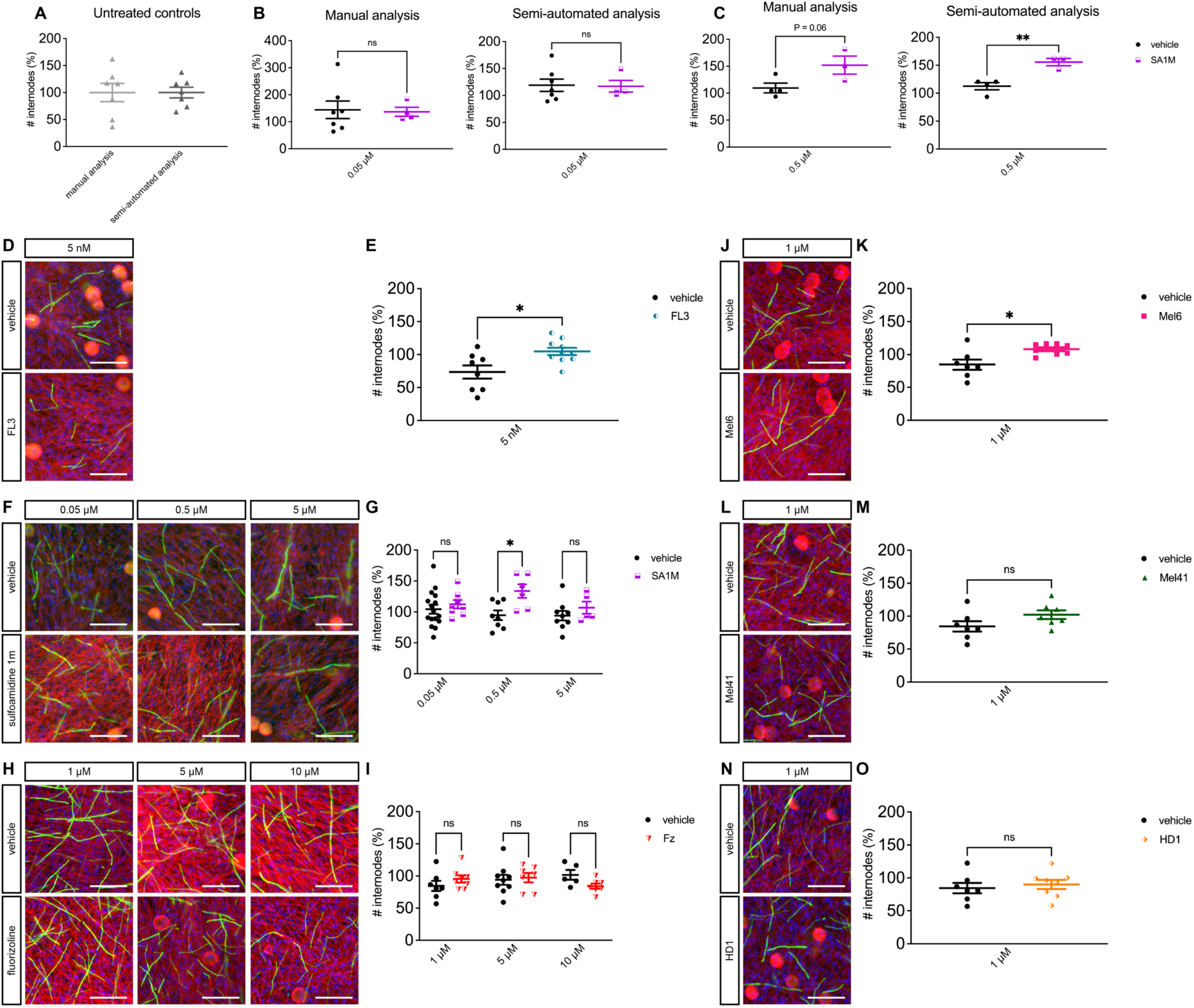
A semi-automated approach for assessing the number of myelinated internodes *in vitro* produces similar results to a manual approach. Lower concentrations of PHB-modulating compounds do not reduce the number of myelinated internodes. (A) Manual or semi-automated analyses produce similar results in the untreated control condition. (B) Manual and semi-automated analyses show that treatment with sulfoamidine 1m (SA1M) at 0.05 μM does not cause a change in the number of myelinated internodes. (C) Manual and semi-automated analyses demonstrate that treatment with SA1M at 0.5 μM increases the number of myelinated internodes. (D-O). Representative immunofluorescent images and corresponding quantifications of the percentage of myelin internodes relative to untreated controls following treatment with six different PHB-modulating compounds, at the concentrations indicated. β-III tubulin immunostaining is shown in red, myelin basic protein (MBP) in green and DAPI is in blue. Scale bar = 100 μm. Each data point represents an individual coverslip, from which the number of internodes was averaged from 3-6 random fields of view. Data were pooled from at least two independent rat preparations of neurons. Data are presented as mean ± SEM and statistical significance as calculated by unpaired t-test, and corrected for multiple comparisons using the Holm-Šídák method where appropriate, indicated as follows: * P < 0.05, ** P ≤ 0.01, *** P ≤ 0.001.

**Supplementary Figure 2.**
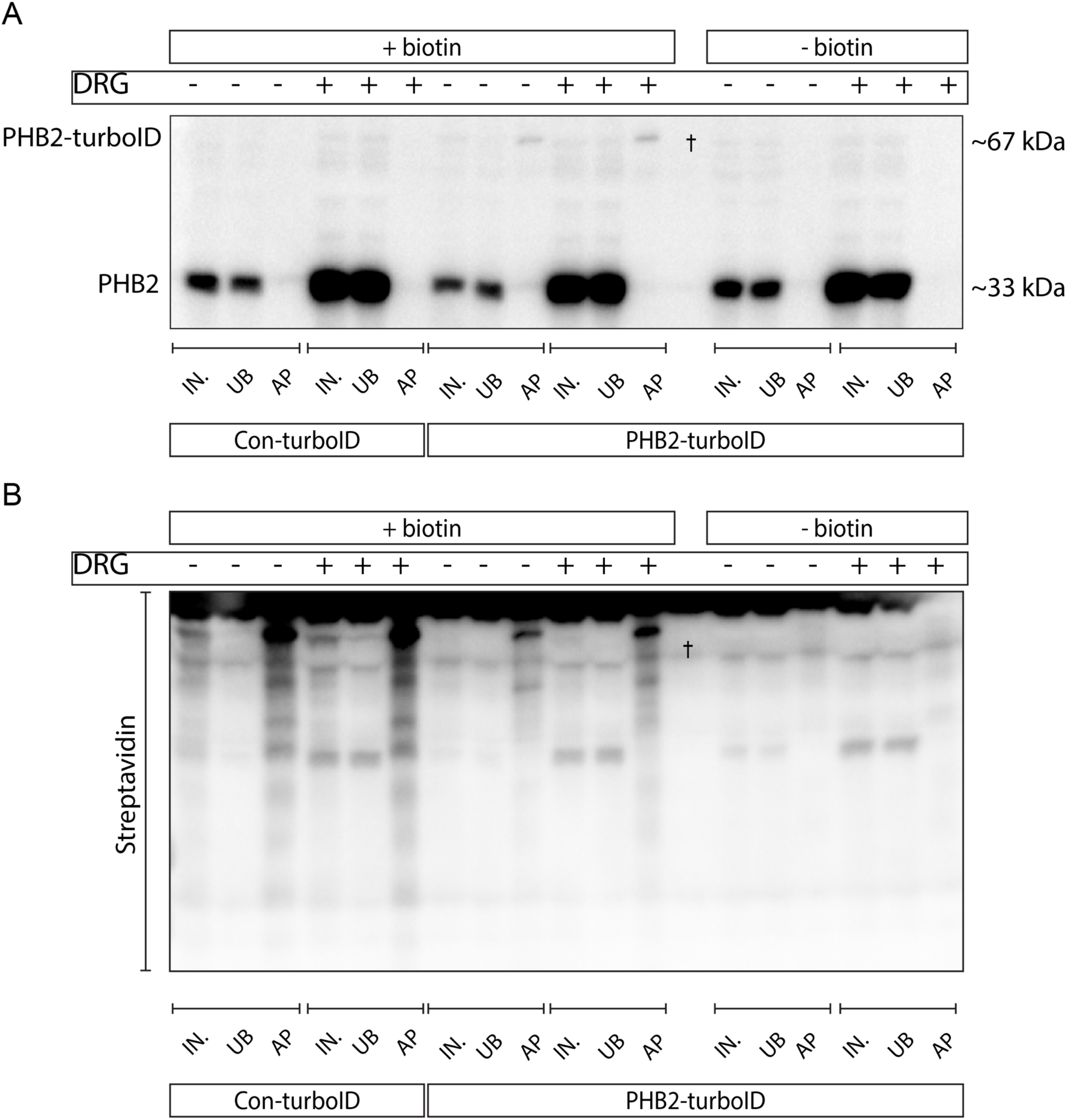
PHB2-turboID is expressed in primary rat Schwann cells and biotinylates proteins. (A) Western blot showing endogenous PHB2 at 33 kDa and a PHB2-immunoreactive band at approximately 67 kDa, corresponding to the size of the PHB2-turboID fusion construct. This band is most clearly evident in the affinity purified (AP) samples in which Schwann cells express PHB2-turboID and are treated with biotin (lanes 9 and 12), suggesting that: 1. PHB2-turboID is expressed at low levels in transfected Schwann cells so that it is only clearly detectable by western blot after affinity purification and 2. PHB2-turboID self-biotinylates. (B) Incubation with streptavidin-HRP reveals that numerous proteins are biotinylated by the PHB2-turboID construct when neurons were both absent and present (lanes 9 and 12, respectively). Biotinylated proteins were not detected when the cells were plated without biotin supplementation (lanes 15 and 18). In contrast and as expected, many more biotinylated proteins were evident in cells expressing the Con-turboID construct (lanes 3 and 6). IN. = input, UB = unbound, AP = affinity purification. ^†^ indicates an empty lane.

**Supplementary Figure 3.**
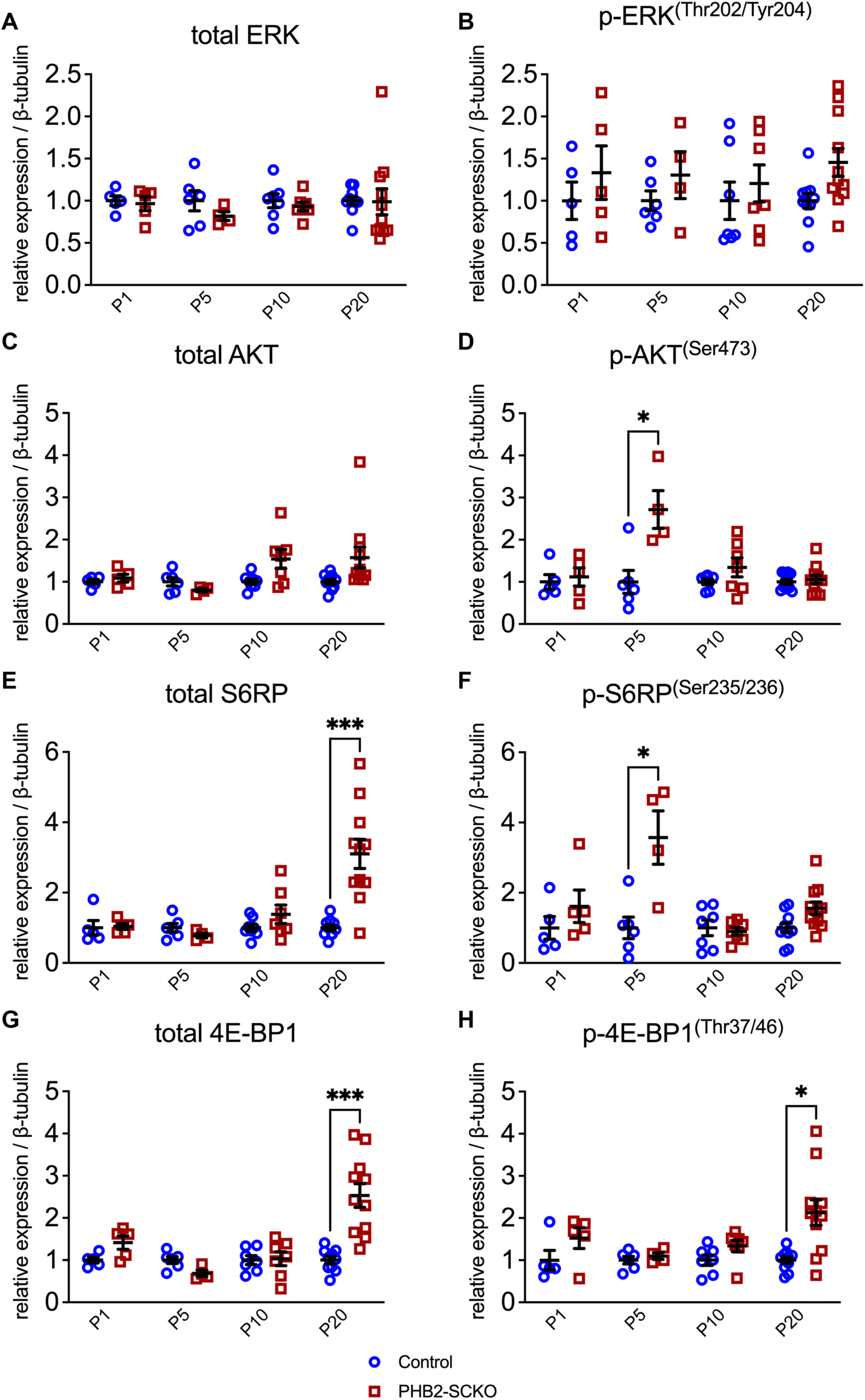
Total and phosphorylated levels of proteins in the ERK, AKT and mTORC1 signaling pathways in control and PHB2-SCKO animals. Quantification of total ERK (A), p-ERK (B), total AKT (C), p-AKT (D), total S6RP (E), p-S6RP (F), total 4E-BP1 (G) and p-4E-BP1 (H) in sciatic nerve lysate from P1, P5, P10 and P20 control and PHB2-SCKO animals, from the western blots shown in Fig. 4 (uncropped blots in Supp. Fig. 7). Each data point represents an individual animal, except for in the P1 and P5 peripheral nerve lysates, when multiple nerves from animals with the same genotype were pooled, and each data point represents an independent pool of nerves. Expression of the protein of interest is calculated relative to the loading control β-tubulin. Expression was then normalised to the average relative expression of the controls, for each age. Data are presented as mean ± SEM and statistical significance, as calculated by unpaired t-test and corrected for multiple comparisons using the Holm-Šídák method, is indicated as follows: * P < 0.05, ** P ≤ 0.01, *** P ≤ 0.001.

**Supplementary Figure 4.**
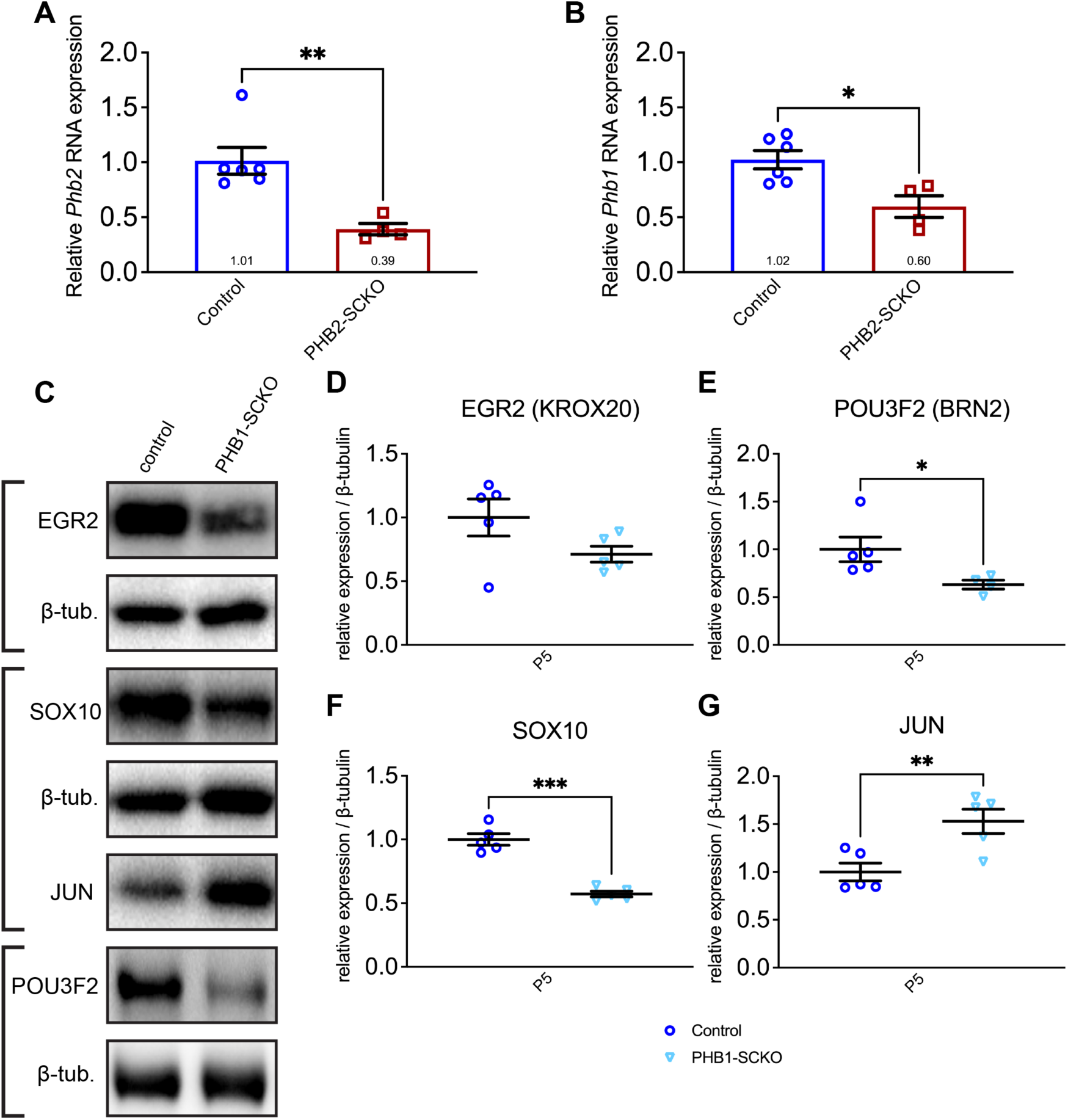
Key Schwann cell transcription factors are also dysregulated in PHB1-SCKO nerves. A reduction in both *Phb2* (A) and *Phb1* (B) mRNA is detected in the peripheral nerves of P20 PHB2-SCKO animals by qRT-PCR. (C) Representative western blots showing EGR2, SOX10, JUN, POU3F2 and β-tubulin (loading control) levels in peripheral nerve lysate from P5 control and PHB1-SCKO animals. (D-G) Quantification of the EGR2 (D), POU3F2 (E), SOX10 (F) and JUN (G) levels from western blots represented in (C). In (A) and (B) each data point represents an individual animal. In (C)-(G) multiple nerves from animals with the same genotype were pooled, and each data point represents an independent pool of nerves. Expression of the protein of interest was calculated relative to the loading control β-tubulin then normalised to the average relative expression of the controls. Data are presented as mean ± SEM and statistical significance, as calculated by unpaired t-test, is indicated as follows: * P < 0.05, ** P ≤ 0.01, *** P ≤ 0.001.

**Supplementary Figure 5.**
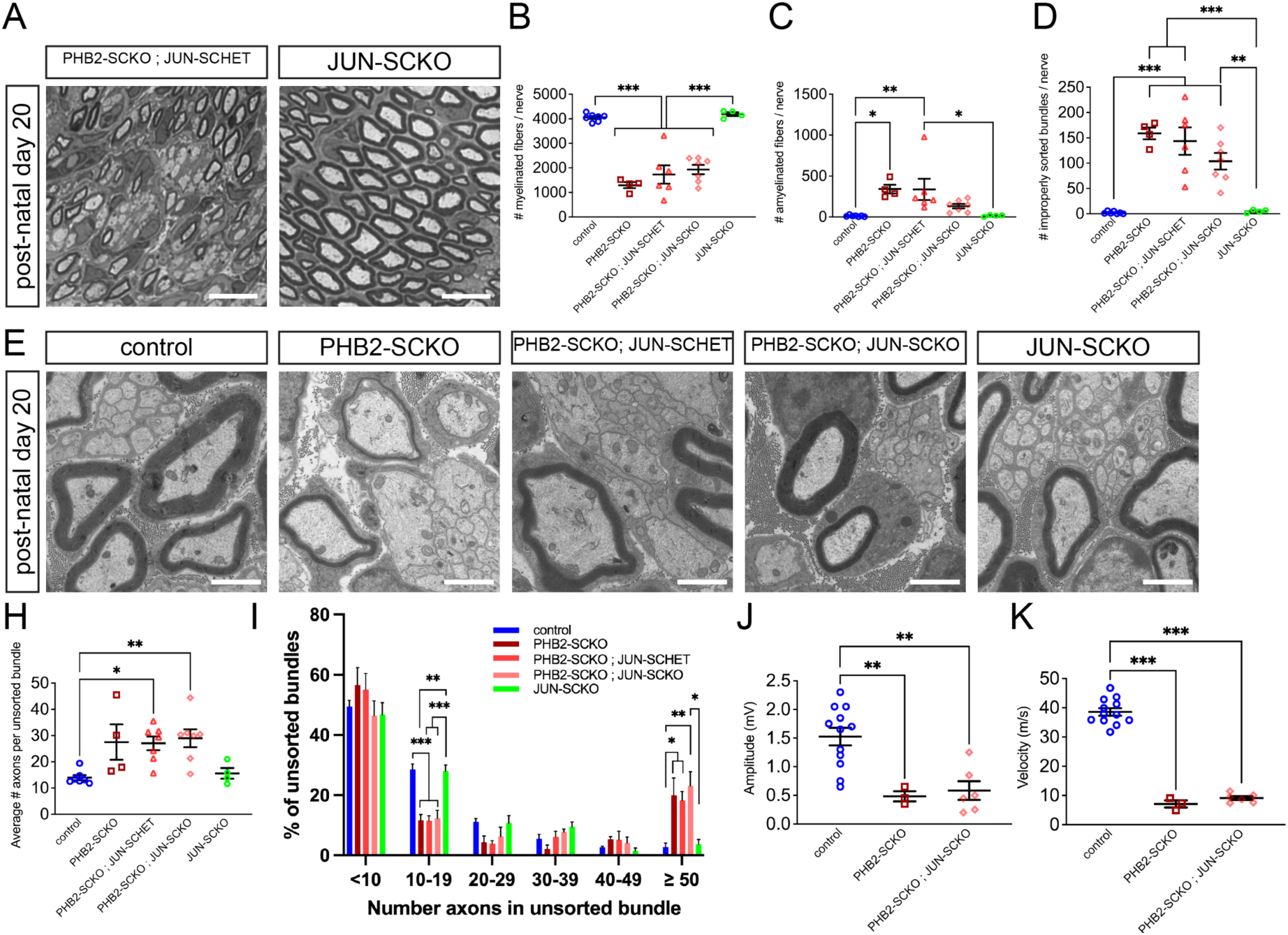
JUN-SCKO confers a partial rescue of the developmental defect in PHB2-SCKO animals. (A) Representative cross sections from P20 sciatic nerves. Scale bar = 10 μm. (B-D) Quantification of the number of myelinated fibers (B), amyelinated fibers (C) and improperly sorted bundles (D) from P20 whole sciatic nerve cross sections represented in (A) and in Fig. 7G. (E) Representative electron micrographs of P20 sciatic nerve cross sections. Scale bar = 2 μm. (H) Quantification of the number of axons per unsorted bundle in P20 sciatic nerves (calculated from electron micrograph images). (I) Distribution of unsorted bundle size by the number of axons in the bundle. (J-K) Compound muscle action potential amplitudes (J) and nerve conduction velocities (K) in P20 mice. Each data point represents an individual animal. Statistical significance was calculated by one-way ANOVA: (B) F (4, 23) = 36.94, P < 0.001, (C) F (4, 23) = 5.556, P = 0.003, (D) F (4, 23) = 19.30, P < 0.001, (H) F (4, 24) = 5.259, P = 0.0035, (I) <10 axons F (4, 24) = 0.9441, P = 0.4556, 10-19 axons F (4, 24) = 17.40, P < 0.0001, 20-29 axons F (4, 24) = 2.650, P = 0.0580, 30-39 axons F (4, 24) = 2.430, P = 0.0753, 40-49 axons F (4, 24) = 0.6615, P = 0.6248, β50 axons F (4, 24) = 7.078, P = 0.0007, (J) F (2, 18) = 11.19, P = 0.0007, (K) F (2, 18) = 180.5, P < 0.0001. Data are presented as mean ± SEM and statistical significance, as determined by Tukey’s multiple comparisons tests, is indicated on the graphs * P < 0.05, ** P ≤ 0.01, *** P ≤ 0.001.

**Supplementary Figure 6.**
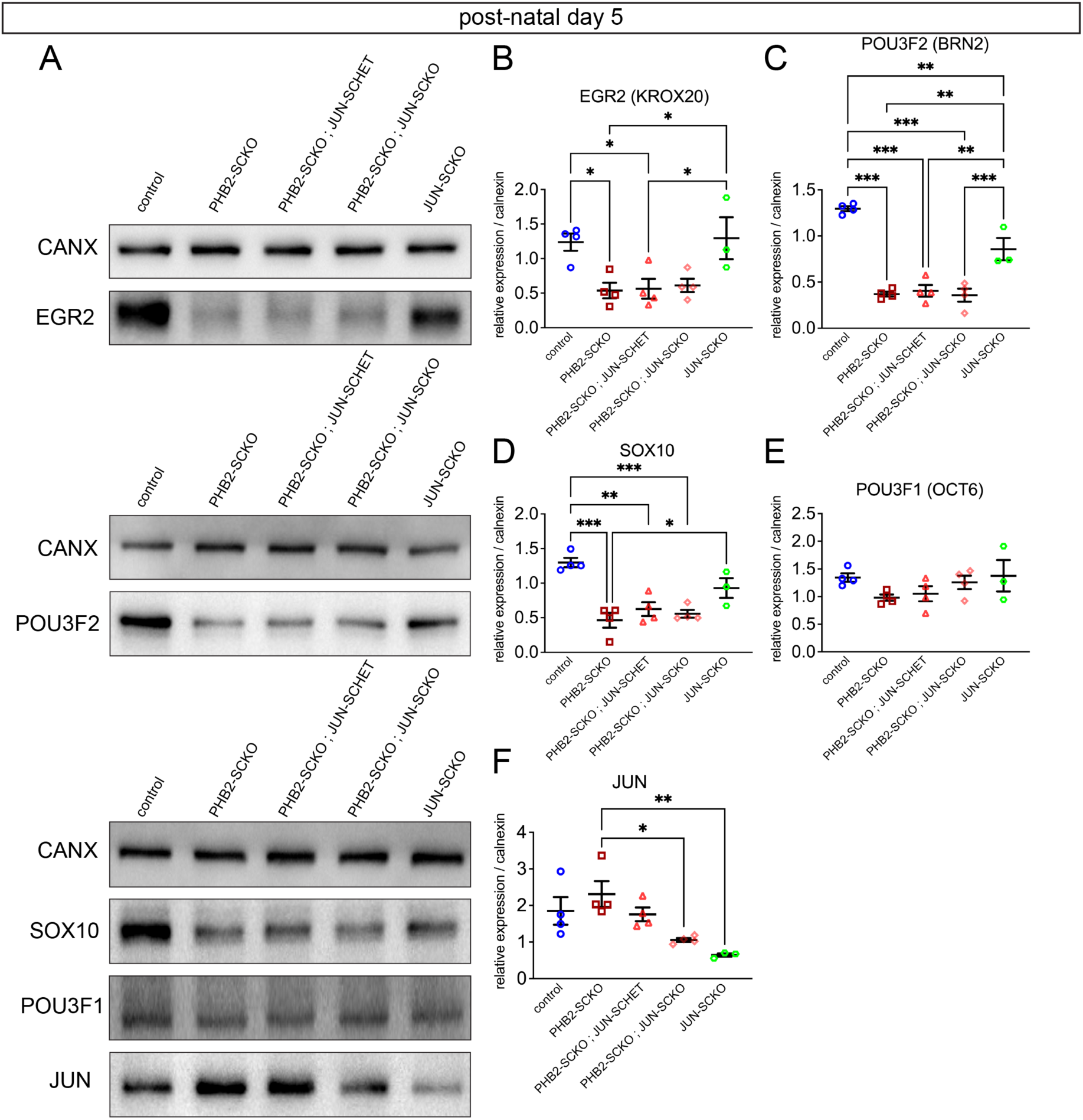
JUN-SCKO does not rescue transcription factor dysregulation in PHB2-SCKO animals. (A) Representative western blots showing EGR2, POU3F2, SOX10, POU3F1, JUN, and calnexin (CANX, loading control) protein levels in peripheral nerve lysate from P5 animals. (B-F) Quantification of the EGR2 (B), POU3F2 (C), SOX10 (D), POU3F1 (E) and JUN (F) levels from western blots represented in (A). Each data point represents an independent pool of nerves. Expression of the protein of interest was calculated relative to the loading control calnexin then normalised to a control reference that was run on every blot. Statistical significance was calculated by one-way ANOVA: (B) F (4, 14) = 5.972, P < 0.0051, (C) F (4, 14) = 44.53, P < 0.0001, (D) F (4, 14) = 13.55, P = 0.0001, (E) F (4, 14) = 1.648, P = 0.2176, (F) F (4, 14) = 6.011, P < 0.0050. Data are presented as mean ± SEM and statistical significance, as determined by Tukey’s multiple comparisons tests, is indicated as follows: * P < 0.05, ** P ≤ 0.01, *** P ≤ 0.001.

**Supplementary Figure 7.**
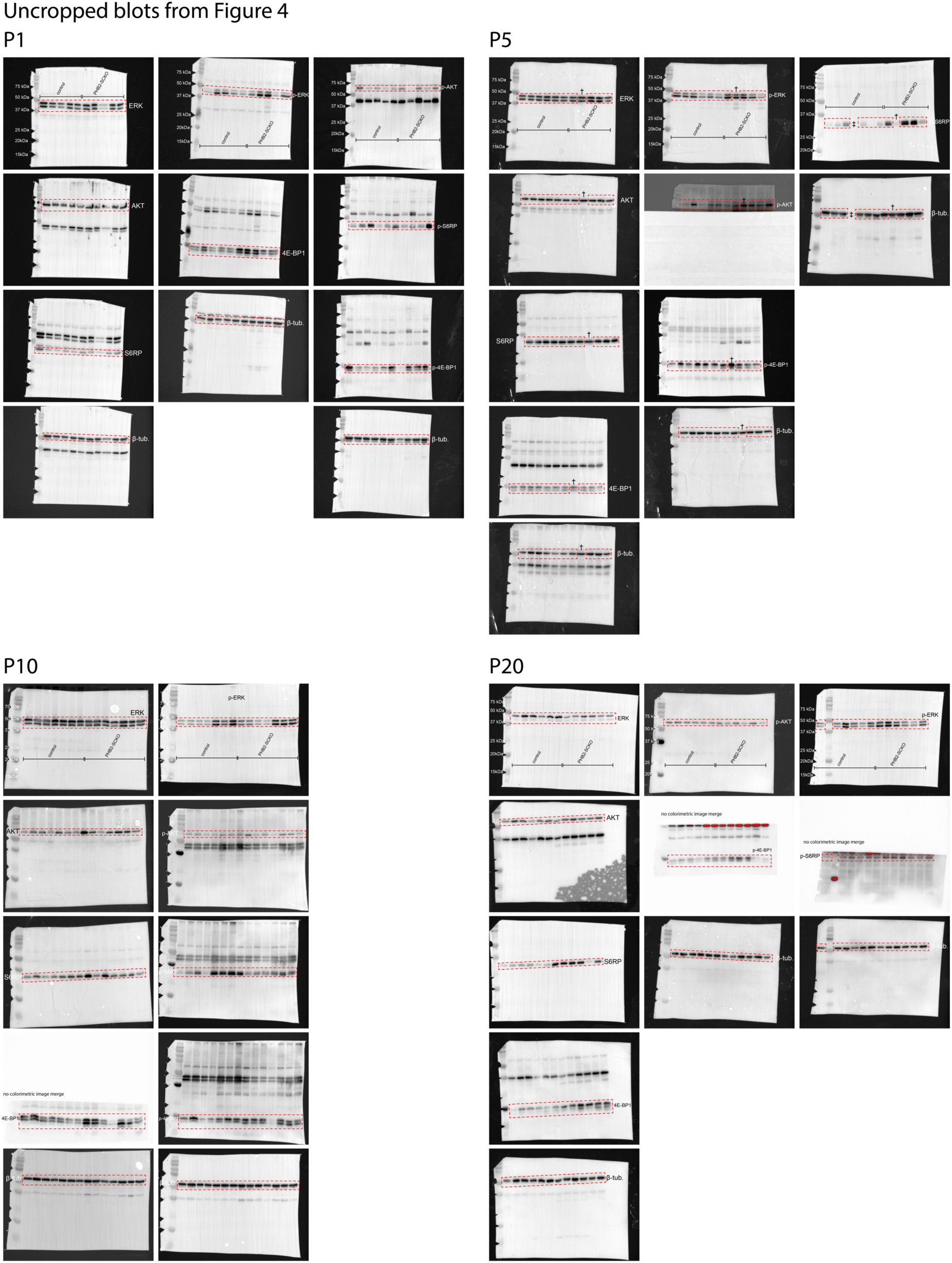
Uncropped blots from Fig. 4. Red dashed lines indicate where the images were cropped for the main figures. Images of the same membrane probed for different proteins of interest are organized vertically. Some membrane images have not been merged with a corresponding colorimetric image (to show the ladder); these have been labeled as such. ^†^ indicates a lysate that was removed from analysis due to genotype uncertainty. ‡ indicates an empty lane.

**Supplementary Figure 8.**
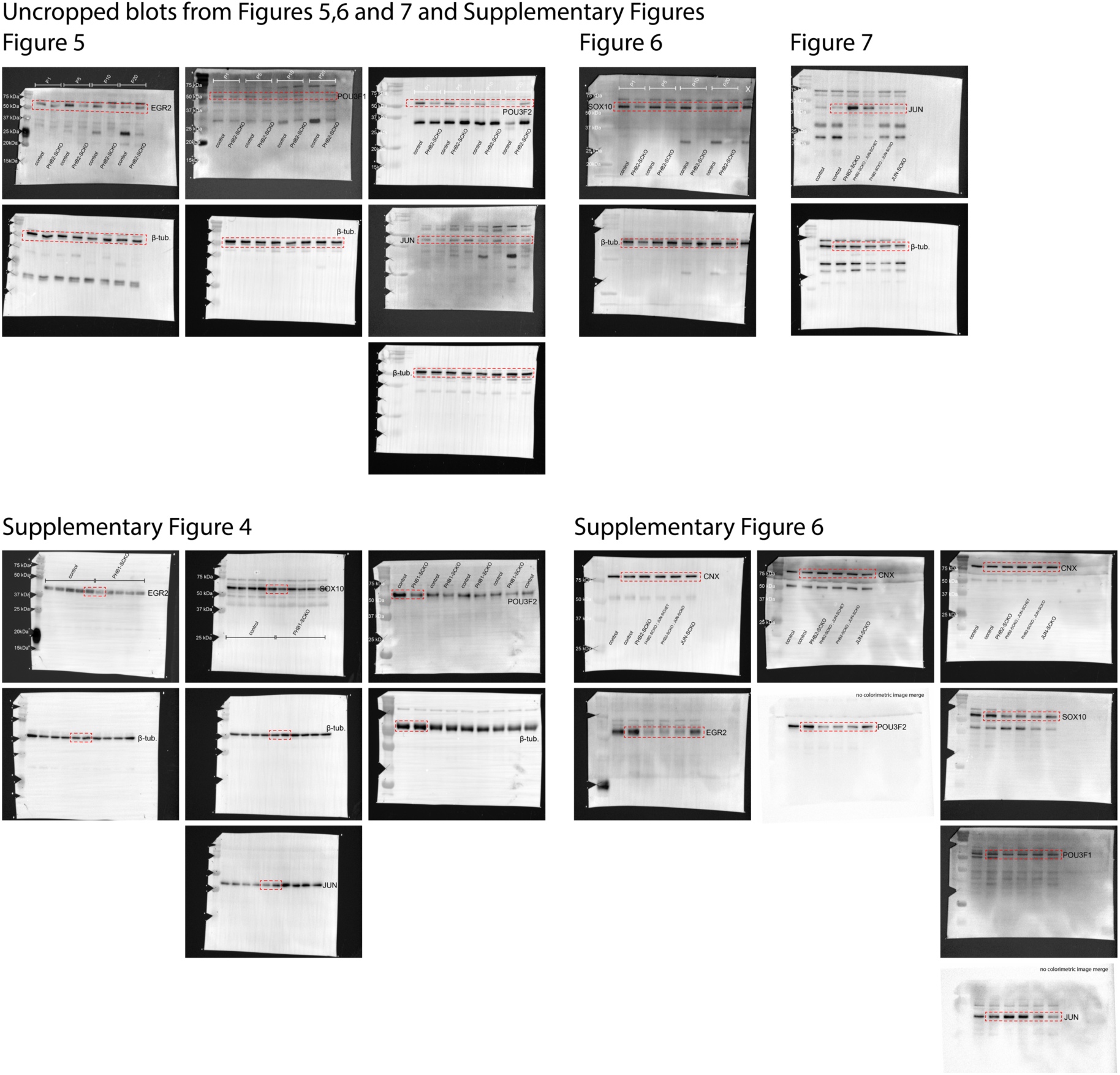
Uncropped blots from Fig. 5, 6 and 7 and Supp. Fig. 4 and 6. Red dashed lines indicate where the images were cropped for the main figures. Images of the same membrane probed for different proteins of interest are organized vertically. Some membrane images have not been merged with a corresponding colorimetric image (to show the ladder); these have been labeled as such.

## References

Arthur-Farraj, P. J., Latouche, M., Wilton, D. K., Quintes, S., Chabrol, E., Banerjee, A., Woodhoo, A., Jenkins, B., Rahman, M., Turmaine, M., Wicher, G. K., Mitter, R., Greensmith, L., Behrens, A., Raivich, G., Mirsky, R., & Jessen, K. R. (2012). c-Jun reprograms Schwann cells of injured nerves to generate a repair cell essential for regeneration. Neuron, 75(4), 633–647. 10.1016/j.neuron.2012.06.021

Arthur-Farraj, P. J., Morgan, C. C., Adamowicz, M., Gomez-Sanchez, J. A., Fazal, S. V., Beucher, A., Razzaghi, B., Mirsky, R., Jessen, K. R., & Aitman, T. J. (2017). Changes in the Coding and Non-coding Transcriptome and DNA Methylome that Define the Schwann Cell Repair Phenotype after Nerve Injury. Cell Reports, 20(11), 2719–2734. 10.1016/j.celrep.2017.08.064

Bacher, S., Achatz, G., Schmitz, M. L., & Lamers, M. C. (2002). Prohibitin and prohibitone are contained in high-molecular weight complexes and interact with alpha-actinin and annexin A2. Biochimie, 84(12), 1207–1220. 10.1016/s0300-9084(02)00027-5

Behrens, A., Sibilia, M., David, J. P., Mohle-Steinlein, U., Tronche, F., Schutz, G., & Wagner, E. F. (2002). Impaired postnatal hepatocyte proliferation and liver regeneration in mice lacking c-jun in the liver. EMBO J, 21(7), 1782–1790. 10.1093/emboj/21.7.1782

Beirowski, B. (2019). The LKB1-AMPK and mTORC1 Metabolic Signaling Networks in Schwann Cells Control Axon Integrity and Myelination: Assembling and upholding nerves by metabolic signaling in Schwann cells. Bioessays, 41(1), e1800075. 10.1002/bies.201800075

Bentayeb, H., Aitamer, M., Petit, B., Dubanet, L., Elderwish, S., Désaubry, L., de Gramont, A., Raymond, E., Olivrie, A., Abraham, J., Jauberteau, M. O., & Troutaud, D. (2019). Prohibitin (PHB) expression is associated with aggressiveness in DLBCL and flavagline-mediated inhibition of cytoplasmic PHB functions induces anti-tumor effects. J Exp Clin Cancer Res, 38(1), 450. 10.1186/s13046-019-1440-4

Blanchard, A. D., Sinanan, A., Parmantier, E., Zwart, R., Broos, L., Meijer, D., Meier, C., Jessen, K. R., & Mirsky, R. (1996). Oct-6 (SCIP/Tst-1) is expressed in Schwann cell precursors, embryonic Schwann cells, and postnatal myelinating Schwann cells: Comparison with Oct-1, Krox-20, and Pax-3. Journal of Neuroscience Research 6(5), 630–640. Doi 10.1002/(Sici)1097-4547(19961201)46:5<630::Aid-Jnr11>3.0.Co;2-0

Boucanova, F., & Chrast, R. (2020). Metabolic Interaction Between Schwann Cells and Axons Under Physiological and Disease Conditions. Front Cell Neurosci, 14, 148. 10.3389/fncel.2020.00148

Branon, T. C., Bosch, J. A., Sanchez, A. D., Udeshi, N. D., Svinkina, T., Carr, S. A., Feldman, J. L., Perrimon, N., & Ting, A. Y. (2018). Efficient proximity labeling in living cells and organisms with TurboID. Nature Biotechnology, 36(9), 880-+. 10.1038/nbt.4201

Britsch, S., Goerich, D. E., Riethmacher, D., Peirano, R. I., Rossner, M., Nave, K. A., Birchmeier, C., & Wegner, M. (2001). The transcription factor Sox10 is a key regulator of peripheral glial development. Genes Dev, 15(1), 66–78. 10.1101/gad.186601

Brockes, J. P., Fields, K. L., & Raff, M. C. (1979). Studies on cultured rat Schwann cells. I. Establishment of purified populations from cultures of peripheral nerve. Brain Res, 165(1), 105–118. 10.1016/0006-8993(79)90048-9

Bunge, R. P., & Bunge, M. B. (1983). Interrelationship between Schwann-Cell Function and Extracellular-Matrix Production. Trends in Neurosciences, 6(12), 499–505. Doi 10.1016/0166-2236(83)90235-7

Della-Flora Nunes, G., Wilson, E. R., Hurley, E., He, B., O’Malley, B. W., Poitelon, Y., Wrabetz, L., & Feltri, M.L. (2021). Activation of mTORC1 and c-Jun by Prohibitin1 loss in Schwann cells may link mitochondrial dysfunction to demyelination. Elife, 10. 10.7554/eLife.66278

Della-Flora Nunes, G., Wilson, E. R., Marziali, L. N., Hurley, E., Silvestri, N., He, B., O’Malley, B. W., Beirowski, B., Poitelon, Y., Wrabetz, L., & Feltri, M. L. (2021). Prohibitin 1 is essential to preserve mitochondria and myelin integrity in Schwann cells. Nat Commun, 12(1), 3285. 10.1038/s41467-021-23552-8

Djehal, A., Krayem, M., Najem, A., Hammoud, H., Cresteil, T., Nebigil, C. G., Wang, D., Yu, P., Bentouhami, E., Ghanem, G. E., & Desaubry, L. (2018). Targeting prohibitin with small molecules to promote melanogenesis and apoptosis in melanoma cells. Eur J Med Chem, 155, 880–888. 10.1016/j.ejmech.2018.06.052

Feltri, M. L., D’Antonio, M., Previtali, S., Fasolini, M., Messing, A., & Wrabetz, L. (1999). P0-Cre Transgenic Mice for Inactivation of Adhesion Molecules in Schwann Cells. Ann N Y Acad Sci, 883(1), 116–123. 10.1111/j.1749-6632.1999.tb08574.x

Feltri, M. L., Poitelon, Y., & Previtali, S. C. (2016). How Schwann Cells Sort Axons: New Concepts. Neuroscientist, 22(3), 252–265. 10.1177/1073858415572361

Figlia, G., Norrmen, C., Pereira, J. A., Gerber, D., & Suter, U. (2017). Dual function of the PI3K-Akt-mTORC1 axis in myelination of the peripheral nervous system. Elife, 6. 10.7554/eLife.29241

Gamble, H. J., & Gosset, J. M. (1966). Specialization of Schwann Cell Membranes. Nature, 212(5063), 734–735. 10.1038/212734a0

Ghislain, J., Desmarquet-Trin-Dinh, C., Jaegle, M., Meijer, D., Charnay, P., & Frain, M. (2002). Characterisation of cis-acting sequences reveals a biphasic, axon-dependent regulation of Krox20 during Schwann cell development. Development, 129(1), 155–166. 10.1242/dev.129.1.155

Goodman, S. R., Johnson, D., Youngentob, S. L., & Kakhniashvili, D. (2019). The Spectrinome: The Interactome of a Scaffold Protein Creating Nuclear and Cytoplasmic Connectivity and Function. Exp Biol Med (Maywood), 244(15), 1273–1302. 10.1177/1535370219867269

Grindheim, A. K., Patil, S. S., Nebigil, C. G., Desaubry, L., & Vedeler, A. (2023). The flavagline FL3 interferes with the association of Annexin A2 with the eIF4F initiation complex and transiently stimulates the translation of annexin A2 mRNA. Front Cell Dev Biol, 11, 1094941. 10.3389/fcell.2023.1094941

He, B., Kim, T. H., Kommagani, R., Feng, Q., Lanz, R. B., Jeong, J. W., DeMayo, F. J., Katzenellenbogen, B. S., Lydon, J. P., & O’Malley, B. W. (2011). Estrogen-regulated prohibitin is required for mouse uterine development and adult function. Endocrinology, 152(3), 1047–1056. 10.1210/en.2010-0732

Jackson, D. N., Alula, K. M., Delgado-Deida, Y., Tabti, R., Turner, K., Wang, X., Venuprasad, K., Souza, R. F., Désaubry, L., & Theiss, A. L. (2020). The Synthetic Small Molecule FL3 Combats Intestinal Tumorigenesis via Axin1-Mediated Inhibition of Wnt/β-Catenin Signaling. Cancer Res, 80(17), 3519–3529. 10.1158/0008-5472.Can-20-0216

Jaegle, M., Ghazvini, M., Mandemakers, W., Piirsoo, M., Driegen, S., Levavasseur, F., Raghoenath, S., Grosveld, F., & Meijer, D. (2003). The POU proteins Brn-2 and Oct-6 share important functions in Schwann cell development. Genes & Development, 17(11), 1380–1391. 10.1101/gad.258203

Jessen, K. R., & Mirsky, R. (2016). The repair Schwann cell and its function in regenerating nerves. J Physiol, 594(13), 3521–3531. 10.1113/JP270874

Kalem, M. C., Subbiah, H., Shen, S., Chen, R., Terry, L., Sun, Y., Qu, J., & Panepinto, J. C. (2022). The interactome of <em>Cryptococcus neoformans</em> Rmt5 reveals multiple regulatory points in fungal cell biology and pathogenesis. bioRxiv, 2022.2001.2013.475903. 10.1101/2022.01.13.475903

Kanagaki, S., Tsutsui, Y., Kobayashi, N., Komine, T., Ito, M., Akasaka, Y., Nagasawa, M., Ide, T., Omae, N., Nakao, K., Rembutsu, M., Iwago, M., Yonezawa, A., Hosokawa, Y., Hosooka, T., Ogawa, W., & Murakami, K. (2023). Activation of AMP-activated protein kinase (AMPK) through inhibiting interaction with prohibitins. iScience, 26(4), 106293. 10.1016/j.isci.2023.106293

Kim, D. I., Birendra, K. C., Zhu, W., Motamedchaboki, K., Doye, V., & Roux, K. J. (2014). Probing nuclear pore complex architecture with proximity-dependent biotinylation. Proc Natl Acad Sci U S A, 111(24), E2453–2461. 10.1073/pnas.1406459111

Kolonin, M. G., Saha, P. K., Chan, L., Pasqualini, R., & Arap, W. (2004). Reversal of obesity by targeted ablation of adipose tissue. Nat Med, 10(6), 625–632. 10.1038/nm1048

Largeot, A., Klapp, V., Viry, E., Gonder, S., Fernandez Botana, I., Blomme, A., Benzarti, M., Pierson, S., Duculty, C., Marttila, P., Wierz, M., Gargiulo, E., Pagano, G., An, N., El Hachem, N., Perez Hernandez, D., Chakraborty, S., Ysebaert, L., Francois, J. H., . . . Moussay, E. (2023). Inhibition of MYC translation through targeting of the newly identified PHB-eIF4F complex as a therapeutic strategy in CLL. Blood, 141(26), 3166–3183. 10.1182/blood.2022017839

Moncunill-Massaguer, C., Saura-Esteller, J., Perez-Perarnau, A., Palmeri, C. M., Nunez-Vazquez, S., Cosialls, A. M., Gonzalez-Girones, D. M., Pomares, H., Korwitz, A., Preciado, S., Albericio, F., Lavilla, R., Pons, G., Langer, T., Iglesias-Serret, D., & Gil, J. (2015). A novel prohibitin-binding compound induces the mitochondrial apoptotic pathway through NOXA and BIM upregulation. Oncotarget, 6(39), 41750–41765. 10.18632/oncotarget.6154

Monk, K. R., Feltri, M. L., & Taveggia, C. (2015). New insights on Schwann cell development. Glia, 63(8), 1376–1393. 10.1002/glia.22852

Nagarajan, R., Svaren, J., Le, N., Araki, T., Watson, M., & Milbrandt, J. (2001). EGR2 mutations in inherited neuropathies dominant-negatively inhibit myelin gene expression. Neuron, 30(2), 355–368. 10.1016/s0896-6273(01)00282-3

Negro, S., Stazi, M., Marchioretto, M., Tebaldi, T., Rodella, U., Duregotti, E., Gerke, V., Quattrone, A., Montecucco, C., Rigoni, M., & Viero, G. (2018). Hydrogen peroxide is a neuronal alarmin that triggers specific RNAs, local translation of Annexin A2, and cytoskeletal remodeling in Schwann cells. Rna, 24(7), 915–925. 10.1261/rna.064816.117

Osman, C., Merkwirth, C., & Langer, T. (2009). Prohibitins and the functional compartmentalization of mitochondrial membranes. J Cell Sci, 122(Pt 21), 3823–3830. 10.1242/jcs.037655

Park, S., Zhao, Y., Yoon, S., Xu, J., Liao, L., Lydon, J., DeMayo, F., O’Malley, B. W., & Katzenellenbogen, B. S. (2011). Repressor of estrogen receptor activity (REA) is essential for mammary gland morphogenesis and functional activities: studies in conditional knockout mice. Endocrinology, 152(11), 4336–4349. 10.1210/en.2011-1100

Parkinson, D. B., Bhaskaran, A., Arthur-Farraj, P., Noon, L. A., Woodhoo, A., Lloyd, A. C., Feltri, M. L., Wrabetz, L., Behrens, A., Mirsky, R., & Jessen, K. R. (2008). c-Jun is a negative regulator of myelination. Journal of Cell Biology, 181(4), 625–637. 10.1083/jcb.200803013

Ploeger, C., Huth, T., Sugiyanto, R. N., Pusch, S., Goeppert, B., Singer, S., Tabti, R., Hausser, I., Schirmacher, P., Désaubry, L., & Roessler, S. (2020). Prohibitin, STAT3 and SH2D4A physically and functionally interact in tumor cell mitochondria. Cell Death Dis, 11(11), 1023. 10.1038/s41419-020-03220-3

Poitelon, Y., Bogni, S., Matafora, V., Della-Flora Nunes, G., Hurley, E., Ghidinelli, M., Katzenellenbogen, B. S., Taveggia, C., Silvestri, N., Bachi, A., Sannino, A., Wrabetz, L., & Feltri, M. L. (2015). Spatial mapping of juxtacrine axo-glial interactions identifies novel molecules in peripheral myelination. Nat Commun, 6, 8303. 10.1038/ncomms9303

Poitelon, Y., & Feltri, M. L. (2018). The Pseudopod System for Axon-Glia Interactions: Stimulation and Isolation of Schwann Cell Protrusions that Form in Response to Axonal Membranes. Methods Mol Biol, 1739, 233–253. 10.1007/978-1-4939-7649-2_15

Qureshi, R., Yildirim, O., Gasser, A., Basmadjian, C., Zhao, Q., Wilmet, J. P., Désaubry, L., & Nebigil, C. G. (2015). FL3, a Synthetic Flavagline and Ligand of Prohibitins, Protects Cardiomyocytes via STAT3 from Doxorubicin Toxicity. PLoS One, 10(11), e0141826. 10.1371/journal.pone.0141826

Rueden, C. T., Schindelin, J., Hiner, M. C., DeZonia, B. E., Walter, A. E., Arena, E. T., & Eliceiri, K. W. (2017). ImageJ2: ImageJ for the next generation of scientific image data. BMC Bioinformatics, 18(1), 529. 10.1186/s12859-017-1934-z

Salameh, A., Daquinag, A. C., Staquicini, D. I., An, Z., Hajjar, K. A., Pasqualini, R., Arap, W., & Kolonin, M. G. (2016). Prohibitin/annexin 2 interaction regulates fatty acid transport in adipose tissue. JCI Insight, 1(10). 10.1172/jci.insight.86351

Salim, C., Boxberg, Y. V., Alterio, J., Fereol, S., & Nothias, F. (2009). The giant protein AHNAK involved in morphogenesis and laminin substrate adhesion of myelinating Schwann cells. Glia, 57(5), 535–549. 10.1002/glia.20782

Sato, T., Komine, T., Renbutsu, M., & Kobayashi, N. (2012). *Novel hydantoin derivative and medicinal agent comprising same as active ingredient* (Japan JP2013230986A Patent No. WO2012026495A1). https://patents.google.com/patent/WO2012026495A1/en?oq=WO%2f2012%2f026495%2c+AMPK+activators

Schindelin, J., Arganda-Carreras, I., Frise, E., Kaynig, V., Longair, M., Pietzsch, T., Preibisch, S., Rueden, C., Saalfeld, S., Schmid, B., Tinevez, J. Y., White, D. J., Hartenstein, V., Eliceiri, K., Tomancak, P., & Cardona, A. (2012). Fiji: an open-source platform for biological-image analysis. *Nature Methods*, *9*(7), 676-682. 10.1038/nmeth.2019

Susuki, K., Raphael, A. R., Ogawa, Y., Stankewich, M. C., Peles, E., Talbot, W. S., & Rasband, M. N. (2011). Schwann cell spectrins modulate peripheral nerve myelination. Proc Natl Acad Sci U S A, 108(19), 8009–8014. 10.1073/pnas.1019600108

Tabti, R., Lamoureux, F., Charrier, C., Ory, B., Heymann, D., Bentouhami, E., & Désaubry, L. (2021). Development of prohibitin ligands against osteoporosis. Eur J Med Chem, 210, 112961. 10.1016/j.ejmech.2020.112961

Takagi, H., Moyama, C., Taniguchi, K., Ando, K., Matsuda, R., Ando, S., Ii, H., Kageyama, S., Kawauchi, A., Chouha, N., Désaubry, L., & Nakata, S. (2022). Fluorizoline Blocks the Interaction between Prohibitin-2 and γ-Glutamylcyclotransferase and Induces p21(Waf1/Cip1) Expression in MCF7 Breast Cancer Cells. Mol Pharmacol, 101(2), 78–86. 10.1124/molpharm.121.000334

Taveggia, C., & Bolino, A. (2018). DRG Neuron/Schwann Cells Myelinating Cocultures. Methods Mol Biol, 1791, 115–129. 10.1007/978-1-4939-7862-5_9

Walko, G., Wogenstein, K. L., Winter, L., Fischer, I., Feltri, M. L., & Wiche, G. (2013). Stabilization of the dystroglycan complex in Cajal bands of myelinating Schwann cells through plectin-mediated anchorage to vimentin filaments. Glia, 61(8), 1274–1287. 10.1002/glia.22514

Wang, D., Tabti, R., Elderwish, S., Abou-Hamdan, H., Djehal, A., Yu, P., Yurugi, H., Rajalingam, K., Nebigil, C. G., & Desaubry, L. (2020). Prohibitin ligands: a growing armamentarium to tackle cancers, osteoporosis, inflammatory, cardiac and neurological diseases. Cell Mol Life Sci, 77(18), 3525–3546. 10.1007/s00018-020-03475-1

Wei, Y., Chiang, W. C., Sumpter, R., Jr., Mishra, P., & Levine, B. (2017). Prohibitin 2 Is an Inner Mitochondrial Membrane Mitophagy Receptor. Cell, 168(1-2), 224–238 e210. 10.1016/j.cell.2016.11.042

Williams, R. L., & Urbe, S. (2007). The emerging shape of the ESCRT machinery. Nat Rev Mol Cell Biol, 8(5), 355–368. 10.1038/nrm2162

Wilson, E. R., Della-Flora Nunes, G., Weaver, M. R., Frick, L. R., & Feltri, M. L. (2020). Schwann cell interactions during the development of the peripheral nervous system. Dev Neurobiol. 10.1002/dneu.22744

Wintachai, P., Thuaud, F., Basmadjian, C., Roytrakul, S., Ubol, S., Desaubry, L., & Smith, D. R. (2015). Assessment of flavaglines as potential chikungunya virus entry inhibitors. Microbiol Immunol, 59(3), 129–141. 10.1111/1348-0421.12230

Yan, C., Gong, L., Chen, L., Xu, M., Abou-Hamdan, H., Tang, M., Désaubry, L., & Song, Z. (2020). PHB2 (prohibitin 2) promotes PINK1-PRKN/Parkin-dependent mitophagy by the PARL-PGAM5-PINK1 axis. Autophagy, 16(3), 419–434. 10.1080/15548627.2019.1628520

Yuan, G., Chen, X., Liu, Z., Wei, W., Shu, Q., Abou-Hamdan, H., Jiang, L., Li, X., Chen, R., Désaubry, L., Zhou, F., & Xie, D. (2018). Flavagline analog FL3 induces cell cycle arrest in urothelial carcinoma cell of the bladder by inhibiting the Akt/PHB interaction to activate the GADD45α pathway. J Exp Clin Cancer Res, 37(1), 21. 10.1186/s13046-018-0695-5

Yurugi, H., Marini, F., Weber, C., David, K., Zhao, Q., Binder, H., Desaubry, L., & Rajalingam, K. (2017). Targeting prohibitins with chemical ligands inhibits KRAS-mediated lung tumours. Oncogene, 36(42), 5914. 10.1038/onc.2017.307

Zhu, B., Zhai, J., Zhu, H., & Kyprianou, N. (2010). Prohibitin regulates TGF-beta induced apoptosis as a downstream effector of Smad-dependent and -independent signaling. Prostate, 70(1), 17–26. 10.1002/pros.21033

